# A Unifying Mechanism for Shared Splicing Aberrations in Splicing Factor Mutant Cancers

**DOI:** 10.1101/2025.10.08.679601

**Authors:** Prajwal C. Boddu, Rahul Roy, Stephen Hutter, Francis Baumgartner, Wenxue Li, Gabriele Todisco, Francesca Ficara, Matteo Giovanni Della Porta, Yansheng Liu, Torsten Haferlach, Manoj M. Pillai

## Abstract

Cancer-associated splicing factor (SF) mutations in *SF3B1*, *U2AF1*, and *SRSF2* induce distinct changes in alternative splicing (AS). Yet these mutations are strikingly mutually exclusive, pointing to a convergent downstream mechanism. We hypothesized this would be reflected in the AS transcriptome. By analyzing transcriptomes of 395 patients with clonal myeloid disorders and 64 healthy donors, we found most AS alterations to be mutation-specific. However, a robust subset, enriched in the retained intron (RI) program, was shared across mutants. These RI events were bidirectional but highly concordant, and mirrored the effects of SRSF1 loss. SF-mutant states induced hypophosphorylation of RS domains in SRSF1, reducing its function. This arose from an altered AMPKα-AKT balance impairing the AKT-SRPK1-SRSF1 axis. A common upstream trigger was activation of DNA damage response (DDR) by transcriptional R-loops, which increased AMPKα signaling and reduced AKT activity. Pharmacologic DDR activation recapitulated reduced AKT/SRPK1 activity and SRSF1 hypophosphorylation, while relieving DDR restored SRSF1 phosphorylation and corrected RI defects. Thus, beyond cis-acting, mutation-specific changes, SF-mutant cancers share a trans-acting, stress-driven AS signature wherein DDR signaling rewires SRSF1 activity impacting AS. Our results link replication stress, kinase signaling, and RNA processing across genetically diverse clonal states, highlighting potential therapeutic approaches at these nodes.

**Highlights:**

- While most splicing changes differ by splicing factor (SF) mutation, certain retained introns are common across subtypes.
- Changes in RI are bidirectional, concordant across mutant groups, and mirrors SRSF1 loss.
- SF mutations activate DDR, triggering an AMPKα/AKT imbalance that culminates in SRSF1 hypophosphorylation.
- Relieving R-loop induced DDR restores SRSF1 phosphorylation and reverses RI.

## Introduction

Mutations in RNA splicing factors (SFs) are common drivers of many clonal states, most commonly in clonal myeloid disorders such as myelodysplastic syndromes (MDS). Among over 300 proteins in the spliceosome machinery, only a handful are commonly mutated: *SF3B1*, *U2AF1*, *SRSF2* and *ZRSR2*.^1,11–16^ Mutations are typically hemizygous (affecting only one allele), hotspot-specific, and non-synonymous (resulting in amino acid changes)^1^ pointing to neo-morphic physiological effects. Analyses of large MDS patient datasets have revealed that, unlike epigenetic mutations which co-occur amongst themselves and with SF mutations, SF-mutations are seldom found together in the same patient (**Fig. S1A**). Such mutual exclusivity suggests negative epistasis, where activation of same or redundant pathways leads to excessive cellular stress.^2^ Thus, SF-mutant states likely converge on shared biological pathways, though the mechanisms underlying their mutual exclusivity remain unclear. One such could be replicative stress and DNA damage response (DDR), arising from R-loops (structures of two DNA and one RNA strands found at areas of active transcription)^3–5^ Our group and others have recently shown that transcription defects underlie such R-loops and replicative stress in *SF3B1, U2AF1*, and *SRSF2* mutant cells.^4^ How such global aberrations in transcription kinetics, replicative stress, and DDR affect RNA metabolism, including AS, remain poorly characterized.

Extensive analyses of mature transcriptomes of patient samples, cell lines and *in vivo* animal models have helped define the impact of SF mutations on the AS program.^6–10^ These studies relied on inferring relative isoform abundance from short read sequencing, which has many limitations, particularly in quantifying some AS changes, like retained introns (RI),^11^ while exonic changes are detected more reliably. Results in general agreement that most splicing changes are modest ( PSI less than 0.1) and largely distinct and non-overlapping within each mutation type. Specific splicing patterns are also associated with each mutation type: *SF3B1* mutations promote the use of novel or cryptic 3’ splice sites,^14,17^ *SRSF2* mutants alter binding preferences from GGNG to CCNG motifs in exons,^3,18^ and *U2AF1* mutants favor non-consensus 3’ splice sites.^19^ Notably, few prior studies^12^ ^13^ have reported an overlap in a subset of AS events-particularly those involving intron retention. Since each of the SFs act at distinct steps of the spliceosome, such a shared signature is unlikely to arise from mutation-specific, cis biochemical effects and is more parsimoniously explained by a unified, trans-acting mechanism.

RNA-binding proteins (RBPs) regulate all aspects of RNA processing including AS.^14^ ^15^ Among these, SR proteins and hnRNPs are the most extensively characterized. Generally, they are functionally antagonistic and balance of their activities serves as a critical determinant of splicing outcomes.^16,17^ The expression, activity, and subcellular localization of these regulatory RBPs are tightly controlled by their phosphorylation status, which directly influences their ability to regulate AS.^18^ ^19^ RBP phosphorylation is orchestrated by both specialized splicing factor kinases, including SRPK1 and CLK1, as well as broader regulatory kinase families such as AKT, AMPKα, and MAPKs, which phosphorylate a wide range of protein substrates involved in many cellular processes.^20^ These kinases are themselves regulated by intrinsic and extrinsic cellular stress signals.^21^ ^22^

In this study, we investigated the mechanisms by which SF mutations lead to a shared pattern of AS. We hypothesized that their shared pattern points to common mechanisms shared across SF mutations. We demonstrate that these AS patterns closely resemble those observed in SRSF1 knockdown or hypophosphorylation, implicating a disruption of SRSF1 activity as a key contributor. SRSF1 dysfunction in SF mutations arises from perturbations in the AKT-SRPK1 signaling axis, coupled with activation of the AMPKα pathway. Notably, these signaling alterations were recapitulated upon inducing DDR in healthy cells and reversed by alleviating R-loop mediated DDR, using RNaseH1 overexpression in SF-mutant cells. Collectively, our findings position SF-mutant MDS as a disease characterized by AMPKα hyperactivation and AKT inhibition, offering potential avenues for therapeutic intervention targeting these critical signaling pathways.

## RESULTS

### Alterations in the retained intron program distinguish SF mutants from healthy bone marrow

To determine common patterns of alternative splicing (AS) across SF-mutations, we analyzed RNA-seq datasets of CD34+ hematopoietic stem and progenitor cells (HSPC) isolated from bone marrow,^13^ of MDS samples with common SF mutations (*SF3B1*, *SRSF2*, *U2AF1*, *ZRSR2)*. This sample cohort includes 227 *SF3B1* mutant, 106 *SRSF2* mutant, 47 *U2AF1* mutant, 15 *ZRSR2* mutant, and 64 wild type/healthy volunteer samples (**Supplementary excel 1**). Consistent with previous reports, the SF mutations did not co-occur in any of the samples, but were found to overlap with driver mutations of epigenetic modifiers (EMs) such as *ASXL1, DNMT3A, TET2,* and *EZH2.*^23^ ^24^ ^25^ ^26–32^ Given this overlap, we further stratified each SF mutation to those with and without co-existing EM mutations (**Fig. 1A**). We estimated AS pattern in these samples using SUPPA2^33^ which relies on transcript isoform abundances estimated using the Salmon^34^ pseudo-alignment tool to quantify the percent spliced-in (PSI) of each AS event. While junction-based methods such as rMATS^35^ also compute PSI and report group-level averages, they are primarily optimized for event-level differential analyses between two groups. In contrast, SUPPA2 generates per-sample PSI values for all splicing events, generating a continuous numerical matrix which can enables downstream transcriptome-wide comparisons and clustering analyses, across the diverse SF mutant and WT samples.

**Figure 1:**
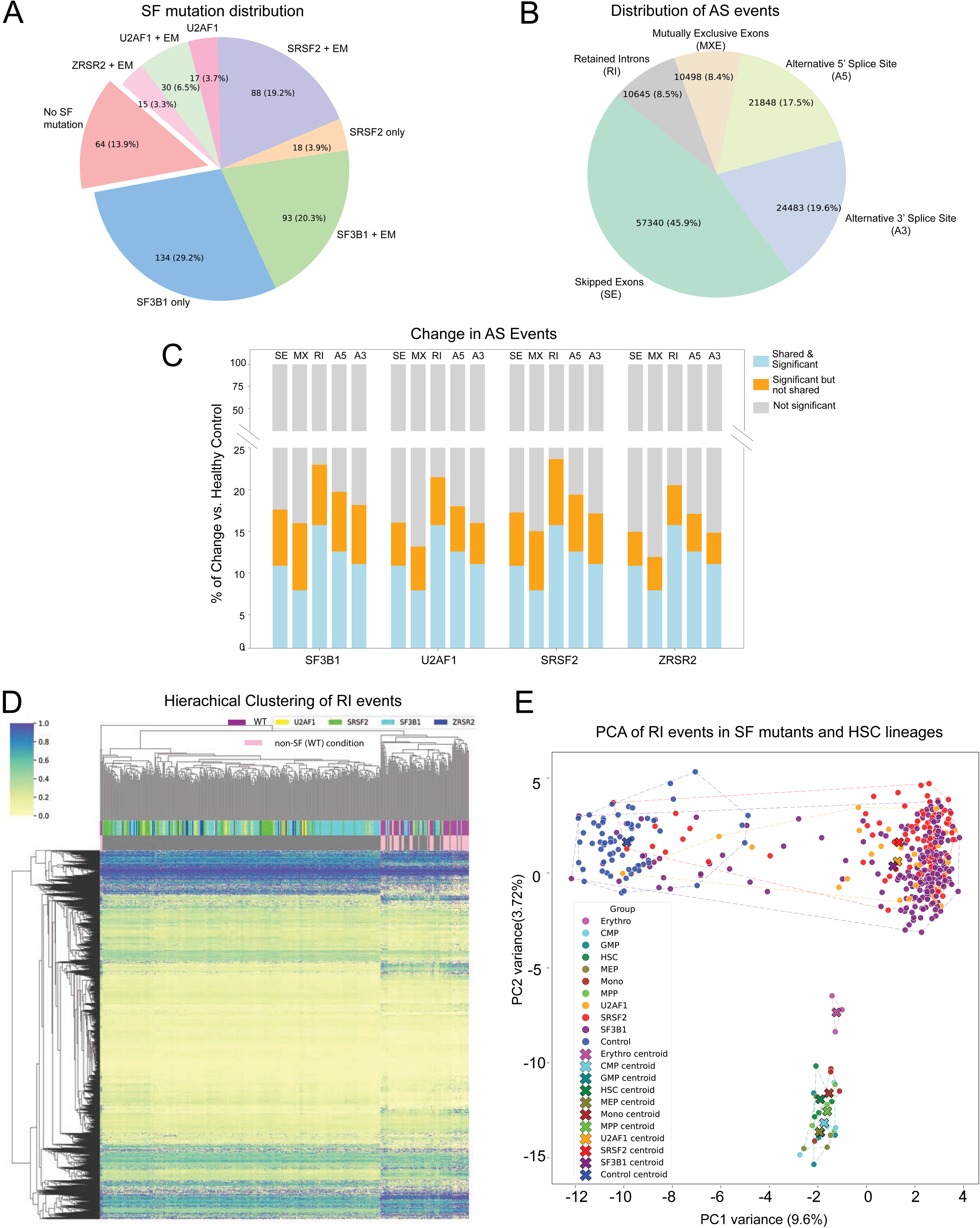
Overview of alternative splicing alterations in SF-mutant MDS. (A) Distribution of splicing factor mutations with or without an epigenetic modifier (EM) co-mutation among 459 samples. Each slice represents a distinct subset. WT indicates healthy volunteer samples. (B) Distribution of AS types across 124815 events by SUPPA2. (C) Stacked bar plot showing the distribution of alternative splicing events across the four SF mutant conditions and five AS types, with bars divided into commonly significant events (blue), unique significant events (not significant across all 4 categories; orange), and non-significant events (gray). Percentages indicate the proportion of events within each condition and AS type. (D) Heatmap showing hierarchical clustering of RI splicing event changes across conditions. Columns represent samples grouped by SF mutations (irrespective of co-mutation status) and controls, with annotations indicating specific conditions. (E) PCA of RI events in SF mutant samples and sorted hematopoietic subpopulations. Each point represents a sample colored by group identity. Samples cluster according to both mutational status and hematopoietic lineage. Notably, SF3B1-, SRSF2-, and U2AF1-mutant samples form distinct, mutation-specific clusters, well-separated from wild-type controls and lineage-defined groups. Erythroblast samples form a discrete cluster with a strong lineage-specific RI program. Convex hulls and centroids (X marks) highlight the spatial distribution and central tendency of each group in the PCA space (PC1 and PC2). Abbreviations used: CMP-common myeloid progenitor; GMP-granulocyte-macrophage progenitor; HSC-hematopoietic stem cell; MEP-megakaryocyte erythroid progenitor; MPP-multipotent progenitor; MONO-CD14+ monocyte.

Using SUPPA2, we quantified 124,815 annotated splicing events (Gencode v46) across five AS categories (skipped exon (SE), retained intron (RI), mutually exclusive exon (MXE), alternative 5’ splice site (A5) or 3’ splice site (A3)), with SE events being the most frequent (**Fig. 1B**). Median transcript-per-million (TPM) values for dominant transcripts were comparable across the SF mutant categories (**Fig. S1B**). However, a smaller fraction of dominant transcripts exceeded 5 TPM in SF mutants (5.31-5.99%) than in WT (11.25%), indicating fewer highly abundant transcripts in the SF mutants. Since PSI estimates are unreliable at low transcript abundance, we restricted analyses to events with dominant transcript ≥5 TPM to ensure reliable comparisons between mutants and WT. Using an absolute ΔPSI ≥ 0.1 and false discovery rate (FDR) ≤0.05, we called significant AS events in each AS category for SF mutant versus WT comparisons. Because SUPPA2 infers RI solely from annotated isoforms in the GTF, we complemented it with IRFinder,^36^ which integrates junction and coverage evidence, and excludes events with low/insufficient coverage and overlap with exonic or 5’/3’ UTR features. The intersection of RI events from both tools yielded high-confidence set (4,151 of 10,645 SUPPA2 events, **Fig. S1C**). RI events were the most shared AS events (12.1%) across SF mutant categories, exceeding MXE, SE, A3 and A5 (**Fig. 1C and Fig. S1D-E,** χ^2^ = 253.5; p-value = 1.1×10⁻□³)). Consistent with this distributional difference, RI events were smore likely than other AS event types to be shared across SF mutants (Odds Ratio (OR): 1.6, p = 6×10⁻²□). To investigate common themes in RI splicing patterns, samples with SF mutations were clustered alongside WT samples. Clustering of samples by RI PSI revealed a clear segregation of SF mutant from WT (**Fig. 1D**) cohorts. This pattern was unchanged by co-occurring EM mutations (**Fig. S1F**). We observed a similar segregation in an independent MDS cohort.^12^ This dataset^12^ utilized poly(A) RNA pull-down followed by sequencing (our dataset: rRNA-depleted total RNA), and hence devoid of nascent intermediaries. This methodological difference, combined with the lower average read depth and length, reduced the total number of analyzable RI events to 879. Despite these limitations, SF-mutant samples again clustered apart from WT (**Fig. S1G**).

MDS is characterized by dysplasia and ineffective hematopoiesis with stage-specific differentiation defects.^37^ ^38^ We therefore asked whether the RI program seen in SF-mutant samples represents a mutation-specific splicing program or instead reflects a developmental block within a particular hematologic lineage. To address this, we used published RNA-seq datasets from sorted hematopoietic subpopulations.^39^ Principal component analysis (PCA) showed close clustering of most subpopulations with the exception of erythroblasts, likely reflecting its distinct transcriptome (**Fig. 1E**). Notably, the patient samples formed distinct, mutation-specific clusters, clearly separated from both WT controls and hematopoietic lineages, including erythroblasts. This pattern was preserved in t-SNE analyses (**Fig. S1H**). Collectively, these findings indicate that the RI patterns in SF-mutant cells do not reflect a differentiation arrest, instead show shared novel, SF-specific changes in the splicing program distinct from normal progenitor populations.

### Intron-retention changes in SF-mutant MDS are enriched for nuclear retained introns

Each SF mutation is associated with characteristic splicing defects: *SF3B1* mutations promote use of cryptic 3’ splice sites,^14,17^ *SRSF2* mutants shift binding preference from GGNG to CCNG motifs in exons,^3,18^ and *U2AF1* mutants favor non-consensus 3’ splice sites.^19^ In addition, prior studies noted a loss of intron retention (LoIR) shared across these mutants. Although intron retention can reflect global splicing failure (e.g., with Pladienolide B), SF3B1, SRSF2, and U2AF1 mutations are neomorphic and do not cause widespread failure; the observed IR changes involve a restricted subset of introns. The biochemical basis of this shared IR program is unclear. We hypothesized that the convergence arises indirectly-from mechanisms outside core splicing catalysis that are common to SF-mutant states. Accordingly, we first focused on RI events (IRFinder) that were consistently excluded (spliced out) across all three SF-mutant states: *SF3B1, SRSF2,* and *U2AF1* mutants.

To ensure consistency, we restricted analysis to introns with a median ≥5 junction-spanning reads per sample in each condition. Significant events were called at |ΔPSI| ≥ 0.1 and FDR < 0.05. As reported previously, LoIR events showed substantial overlap across the three SF-mutant groups, with >50% of LoIR shared by each group (**Fig. 2A**). Gain-of-IR (GoIR) events also overlapped across mutants, though to a lesser extent (**Fig. 2B**). Notably, among the 7,003 RI events in the combined LoIR+GoIR superset, none were discordant-no LoIR event in one mutant class appeared as GoIR in another, and vice versa. Thus, RI alterations in SF mutants are largely shared across mutation classes, occur in both directions (LoIR and GoIR), and are highly concordant, supporting a common trans-acting mechanism underlying these changes.

**Figure 2:**
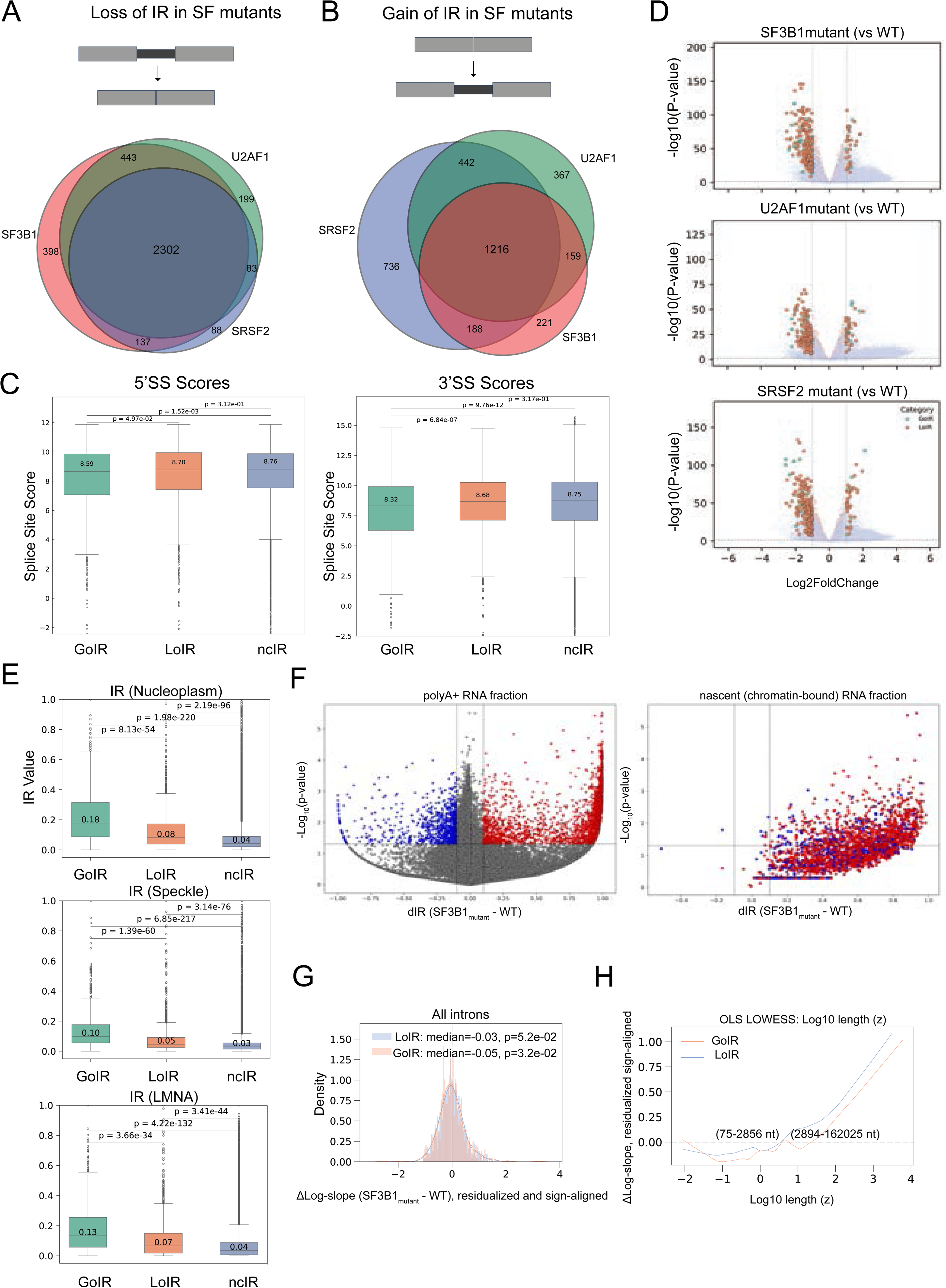
Retention of intron events undergo bidirectional concordant changes across the major SF mutant categories. (A) Venn diagram showing introns with significant loss of intron retention (dRI > 0.1 in SF mutant vs. WT, minimum junction coverage ≥10, Wilcoxon FDR < 0.05). Numbers indicate uniquely or commonly lost RI events across SF3B1, SRSF2, and U2AF1 mutants. (B) Venn diagram showing introns with significant gain of intron retention (dRI < −0.1 in SF mutant vs. WT), under the same filtering criteria as in Fig. 2A. Numbers indicate uniquely or commonly gained RI events across the three SF mutants. (C) Boxplot depicting the distribution of 5’ splice site scores (left panel) and 3’ splice site scores for introns categorized as GoIR, LoIR, and ncIR. The median splice site scores for each category are annotated within the boxplots. Statistical significance between categories was assessed using pairwise comparisons, and p-values are shown. (D) Volcano Plots of Differentially Expressed Genes in Splicing Factor Mutants Compared to Wild-Type (WT), illustrating the log2 fold change of gene expression against the -log10 p-value. Significantly changing genes, associated with introns categorized as GoIR, LoIR, ncIR introns, are highlighted in green, orange, and blue, respectively. (E) Intron Retention (RI) Values for Nuclear sub-compartments Based on HEK293T-APEX-Seq Data. Boxplots represent the baseline RI values for introns categorized as GoIR, LoIR, and ncIR across three nuclear subcompartments: nucleoplasm, Lamina-associated regions (LMNA), and nuclear speckles. Median RI values are annotated within each boxplot. P-values denote pairwise comparisons between categories, indicating significant differences in RI behavior across nuclear subcompartments. (F) Differential RI in SF3B1 Mutant vs. WT across PolyA+ and Chromatin RNA fractions. The volcano plot depicts the changes in RI (dRI) between SF3B1 mutant and WT samples. Top: polyA+ RNA fraction shows both increased and decreased RI events. Bottom: Nascent (chromatin-bound, polyA depleted) RNA displays a consistent increase in RI, indicating primary defects at the co-transcriptional level. Red and blue dots denote the significantly increased or decreased RI in the corresponding polyA+ fraction, respectively (|dRI| > 0.1, p < 0.05). (G) Panels show the distribution of Δlog-slope (SF_mutant_ − WT). Δlog-slope values are residualized vs WT and sign-aligned to WT, so negative values indicate a flatter-than-expected 3′→5′ coverage gradient in the mutant. LoIR (blue) and GoIR (salmon). Shaded histograms with overlaid KDEs are shown; the dashed line marks 0. Text in each panel reports the one-sided Wilcoxon p-value testing median < 0 for each group. (H) Length-dependent shift in slope (OLS LOWESS). LOWESS of the residualized, sign-aligned Δ log-slope (SF_mutant_ − WT) versus z-scored log10 intron length. Dashed line = no change. Numbers in parentheses give the back-transformed length ranges for this dataset: shorter introns (∼75-2,850 nt) tend to show slight flattening (Δ ≤ 0), whereas longer introns (∼2,900-162,025 nt) show progressive steepening (Δ > 0). LoIR (blue) exhibits a stronger length-dependence than GoIR (orange).

To identify features linked to RI changes, we compared introns classified as LoIR, GoIR, or no change (ncIR). At baseline, LoIR and GoIR introns had higher RI than ncIR (median PSI 0.34 and 0.06 vs 0.01). Thus, introns with changes in their retention patterns are not fully spliced out at baseline but are partially retained even before mutation, whereas efficiently spliced introns at baseline are less likely to change with SF mutations. This suggests that LoIR and GoIR have likely sequence features which allow modulation of splicing (increase or decrease) based on additional context, unlike ncIR which are constitutively spliced out. We then examined intron length (**Fig. S2A**), GC content (**Fig. S2B**), splice site strength/score (SSS; a measure of splicing efficiency) at both 5′ and 3′ splice junctions (**Fig. 2C**), and RNA free energy (ΔG) at splice sites (**Fig. S2C**). Both LoIR and GoIR introns were significantly shorter than ncIR introns (**Fig. S2A**), consistent with reports that shorter introns are retention-prone.^40^ GoIR introns also had significantly weaker 5′ and 3′ SSs (**Fig. 2C**) and more negative ΔG scores at splice junctions (**Fig. S2C**). By contrast, 3′ splice site conservation (sc) were similarly low across GoIR, LoIR, and ncIR categories (**Fig. S2D**). Finally, genes harboring GoIR, LoIR, or ncIR introns showed no consistent expression differences (**Fig. 2D**), arguing that RI changes aren’t simply explained by differential gene expression. The similar 3′SS conservation across classes and the lack of concordant gene-expression changes argue against a purely sequence-encoded or transcription-driven model and instead point to trans-acting regulation. Since trans-acting RNA binding proteins (RBPs) such as SR proteins and hnRNPs regulate IR,^41^ ^42^ ^43^ disrupted RBP recruitment in SF mutants may drive the observed changes in RI.

Subcellular localization of RI-RNAs is a critical determinant of its fate. Since our analyses use steady-state RNA-seq from whole cells, we cannot infer whether RI contributes to degradation of mRNA (by exosomes in the nucleus or NMD in cytoplasm). RI-RNAs determined from RNA-seq of total cellular lysates captures both nuclear and cytoplasmic RI-RNAs, with nuclear RI-RNAs thought to predominate.^44^ To assess physiological localization of these RI-RNAs, we leveraged publicly available RNA-seq datasets of nuclear sub compartments (nucleoplasm, speckle, lamina) determined by APEX-labeling and pull down.^45^ Although HEK293T and CD34+ cells have distinct transcriptomes, we reasoned that the sub-compartmental distribution of RNA is conserved for broadly expressed transcripts. LoIR introns were highly enriched for nuclear-retained introns (43%, 22%, and 37% retention in nucleoplasm, speckle, and lamina, respectively, compared to 22%, 10%, and 21% for ncIR (p < 2.2 x 10^-16^ for all comparisons, **Fig. 2E**). GoIR introns exhibited an even higher nuclear retention across sub-compartments, with 70%, 49%, and 60% retention in nucleoplasm, speckle, and lamina, respectively; p < 2.2 x 10^-16^ for all comparisons with ncIR)). Thus, RI events that change in SF-mutant states (LoIR or GoIR) are disproportionately nuclear-retained.^44^ This pattern argues against nonspecific splicing noise and instead implicates a non-random, pathogenic mechanism in which SF mutations perturb a specialized nuclear regulatory layer controlling transcript stability and export.

### A co-transcriptional mechanism underlies RI alterations in SF-mutant MDS

Most introns are removed co-transcriptionally, but a subset is spliced post-transcriptionally. To pinpoint where RI defects arise in SF-mutant cells, we fractionated RNA from two SF3B1-mutant patients and healthy donors into chromatin-associated (nascent, co-transcriptional) and poly(A)+ (nucleoplasmic/cytoplasmic, post-transcriptional) pools.^46^ The chromatin fraction is enriched for nascent pre-mRNAs and reflects co-transcriptional splicing, whereas the poly(A)+ pools reflect post-transcriptional processing. As expected, poly(A)+ RNA showed bidirectional RI changes (GoIR and LoIR) (**Fig. 2F**). Notably, the chromatin fraction consistently exhibited increased RI in mutants, independent of the poly(A)+ direction (**Fig. 2F**). This divergence indicates a primary defect in co-transcriptional splicing (CoTS), with downstream post-transcriptional steps subsequently driving individual introns toward GoIR or LoIR.

As an orthogonal test, and because nascent chromatin RNA-seq was not available for the patient cohort, we leveraged total RNA-seq (which contains both polyadenylated and nascent species) to derive an informatic proxy for CoTS.^47^ In such data, intronic read density typically decays from 5′ to 3′ across an intron;^48^ the magnitude of this 5′→3′ decay (“intron slope”) reflects the balance between Pol II elongation and co-transcriptional splicing (Methods). We summarized this decay with an intronic 5′→3′ slope (reported as a log-ratio slope), which served as a proxy for CoTS efficiency: When CoTS slows, the previously depleted end of the intron gains relatively more unspliced coverage (5′ if the WT profile is 3′-biased; 3′ if it is 5′-biased). Thus, when CoTS slows, intronic coverage becomes more uniform and the slope flattens toward 0. Consistently, across the three SF mutant categories (**Fig 2G, Fig. S2E**), slopes shifted significantly toward flatter values. This flattening occurred in both GoIR and LoIR sets, consistent with a global reduction in CoTS irrespective of steady-state RI outcome. In a multivariable model (adjusting for GC content, ΔG, 5′SS and 3’SS scores and IR class (LoIR vs GoIR)), intron length was the only independent predictor for CoTS slowdown (**Fig. S2F**). Shorter introns showed the largest flattening, whereas long introns showed little to no shift (**Fig. 2H; Fig. S2E, S2G**), supporting a kinetic-coupling model: when co-transcriptional splicing slows, short introns-with a brief window for spliceosome assembly as Pol II passes- are disproportionately affected. Together, these data reconcile the opposite steady-state RI directions: when CoTS falls, introns with limited post-transcriptional capacity (weak signals/branchpoints, detained-intron architecture) tend to accumulate as GoIR, whereas introns with stronger post-transcriptional processing or preferential nuclear clearance complete splicing or are degraded after cleavage/polyadenylation, yielding LoIR despite the CoTS defect.

### Disruption of SRSF1-hnRNP balance driving RI pattern alterations in SF-mutant MDS

We reasoned that, given their distinct binding sites on the pre-mRNA and disparate biochemical effects when mutated, the concordant RI alterations evident in our analysis of MDS samples are not from neomorphic mutant SF function. Instead, we posited a shared downstream trans-acting mechanism. RBPs are primary regulators of AS, including RI. Among the hundreds of RBPs regulating AS, SR and hnRNP proteins are best studied, where they have largely opposing functions.^16,17^ RBP activities are frequently phosphorylation dependent, providing a plausible conduit from SF-mutant stress signaling to trans regulation.^20^ We asked whether SF-mutant RI profiles resemble any of the RBP knockdown (KD) states. To test this, we analyzed RNA-seq datasets of RBP knockdown from the ENCODE study (shRNA knockdowns of 231 RBPs in HepG2 cells and 143 RBPs in K562 cells (listed in **Fig. S3A**)). Using the 3518 RI events with significant dRI (derived from **Fig. 2A-B**) shared across all three SF mutants, we performed unsupervised clustering. SF mutant conditions clustered with various RBP KD conditioning of the erythroleukemia-derived K562 cells (**Fig. S3B**), while the liver-derived HepG2 RBP KD conditions showed much weaker correspondence. We infer that this is likely from similar transcriptomes and splicing programs between these cellular contexts.

We prioritized RBP KD conditions with the strongest positive (closest clustering distance) and negative (most distant clustering) correlations with the SF-mutants (**Fig. 3A**). SRSF1 KD showed the highest concordance with SF-mutant RI changes, whereas hnRNPA0 and hnRNPAB KDs showed the strongest anticorrelation (**Fig. 3B**). These relationships were most pronounced for RI and weaker for other AS classes (SE, MXE, A5/A3; Fig. S3C-D). Together, the data indicate that SF-mutant RI patterns phenocopy SRSF1 loss and oppose hnRNPA0/AB depletion, consistent with a model in which shifts in the SR vs hnRNP balance drive the shared RI signature.

**Figure 3:**
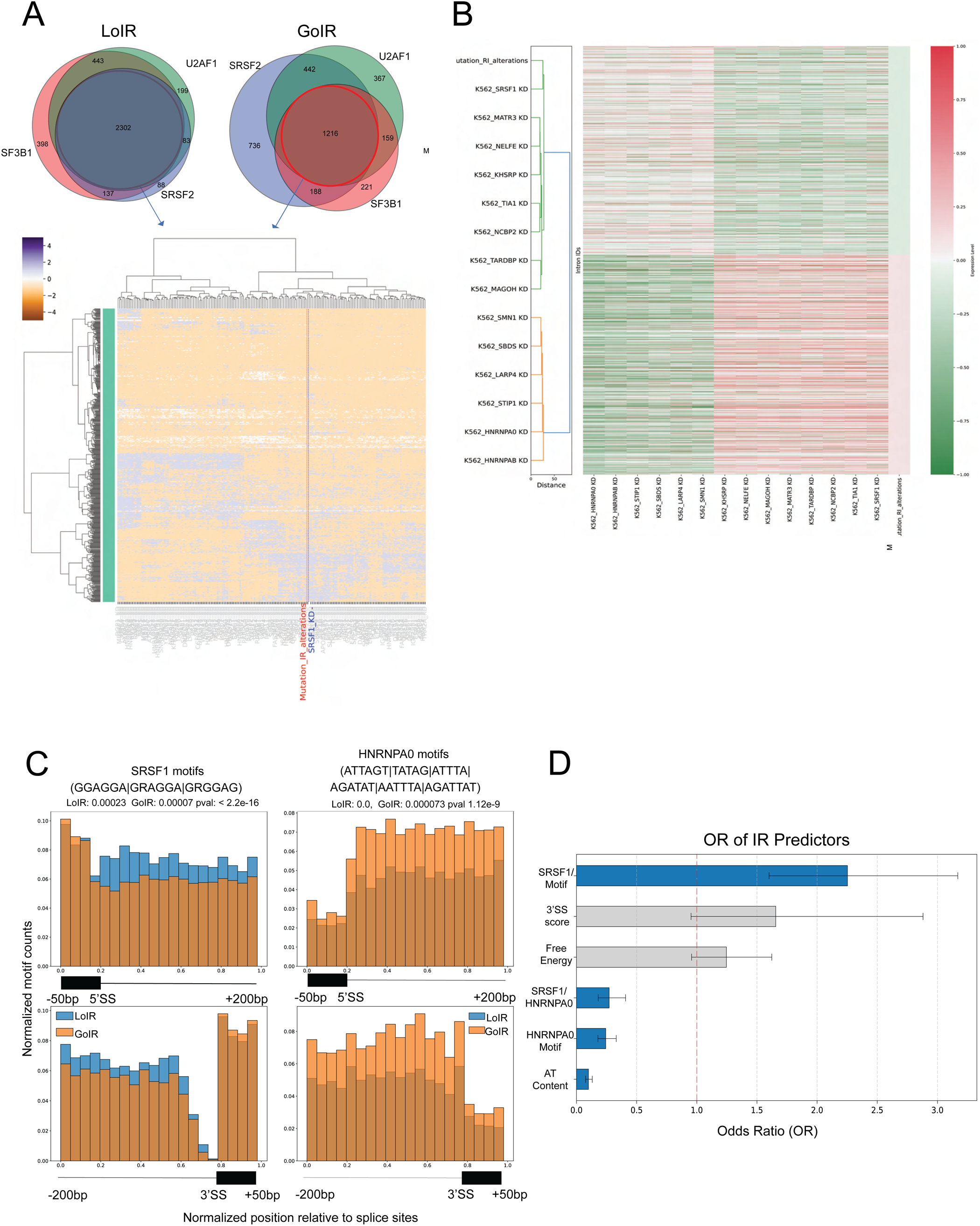
Clustering-based approach to identify key RBPs dysregulated in SF-mutant MDS. (A) Heatmap illustrating the hierarchical clustering of delta RI (dRI) changes involving 3,518 RI events, identified across three splicing factor mutation categories, as detailed in (Fig. 2A-B), across multiple shRNA knockdown conditions in K562 cells. Each row represents an individual RI event, while each column corresponds to a specific knockdown condition. The color intensity reflects the magnitude of dRI changes, with red indicating increased retention and blue indicating decreased retention. Notably, the dendrogram highlights the close clustering of mutation-induced RI alterations and the SRSF1 knockdown condition, emphasized in red and blue, respectively. Clustering was performed on effect directionality; no statistical weights were applied. (B) Focused heatmap illustrating the RI pattern changes, highlighting those that most closely correlate or anticorrelate with mutation-induced RI alterations. Notably, the SRSF1 knockdown condition is emphasized, demonstrating its proximity to mutation-induced RI changes. (D) Histograms depicting the distribution of SRSF1 and hnRNPA0 motif densities within 200 bp intronic regions flanking the splice sites in LoIR_SRSF1KD_ and GoIR_SRSF1KD_ intron categories. The x-axis represents the position relative to the splice site, binned in 10 bp intervals, while the y-axis shows the normalized motif count. Blue bars correspond to LoIR_SRSF1KD_ introns, and orange bars to GoIR_SRSF1KD_ introns. These histograms illustrate the differential motif enrichment patterns near splice sites between the two intron categories. The brown color portions in the bars indicate overlapping regions where both LoIR and GoIR motif densities are present. (D) Logistic regression analysis of intron retention predictors. The forest plot visualizes odds ratios with 95% confidence intervals. Predictors with significant associations (blue bars) include SRSF1 motif density, hnRNPA0 motif density, SRSF1/hnRNPA0 ratio, and AT content.

SRSF1 is a master regulator of AS, which typically promotes splicing of exons by binding to exon splicing enhancer elements. However, its role is highly context-dependent, as it can also inhibit splicing when bound to intronic splicing silencer (ISS) elements.^49^ To better understand the mechanisms driving RI changes upon SRSF1 KD, we examined sequence and motif features distinguishing GoIR_SRSF1KD_ (increased RI in SRSF1 KD; dRI > |0.1|, FDR < 0.05) from LoIR_SRSF1KD_ (decreased RI in SRSF1 KD; dRI > |0.1|, FDR < 0.05). First, we determined the densities of putative SRSF1 and hnRNPA0 binding motifs within 200-bp intronic regions flanking the 5’SS and 3’SS in GoIR_SRSF1KD_ and LoIR_SRSF1KD_ introns. LoIR_SRSF1KD_ introns were enriched for SRSF1 ISS-like motif densities, whereas GoIR_SRSF1KD_ introns carried more HNRNPA0 motifs. and less abundant in LoIR_SRSF1KD_ introns (**Fig. 3C, Fig. S3E**). GoIR_SRSF1KD_ introns also have weaker 3’ SSS (**Fig. S3F**), lower GC content in regions flanking these SSs (**Fig. S3G**), and reduced stability of secondary structures at SSs (**Fig. S3H**). Together, these features suggest that many GoIR introns are poised (weaker baseline splicing) and whose fate depends on the RBP milieu**-**with hnRNPA0 favoring retention and SRSF1 promoting excision-explaining why SRSF1 KD causes GoIR_SRSF1KD_. In contrast, in introns where SRSF1 ISS motifs are enriched, splicing is enhanced upon SRSF1 KD, resulting in reduced RI (LoIR_SRSF1KD)_.

To assess the predictive value of these features in splicing outcomes upon SRSF1 KD, we employed logistic regression with six predictors (SRSF1 motifs, hnRNPA0 motifs, their ratio, AT content, 3′SS strength, free energy). Collinearity was low (variance inflation factor (VIF)^50^ < 5; Methods). hnRNPA0 motif burden, the SRSF1/hnRNPA0 ratio, and AT content were the best predictors (**Fig. 3D**, **Fig. S3I**). Higher AT content significantly lowered the odds of intron loss (OR = 0.10, p < 0.001); greater SRSF1 motif burden increased intron excision, whereas higher hnRNPA0 motif burden predicted retention. Accuracy of the regression model was ∼60% across normalization approaches (Methods), suggesting unmodeled, non-sequence determinants, such as RBP milieu and chromatin state, among others, may be contributing substantially to the outcome. Nonetheless, while not exhaustive, these results support intronic sequence context as a key driver of SRSF1-dependent AS behavior.

### Phosphorylation-dependent changes to SRSF1 function in SF mutant cell lines

SRSF1 phosphorylation is required for nuclear import and for binding to pre-mRNA/RBPs to regulate splicing.^51^ ^52^ ^53^ Conversely, hypophosphorylated SRSF1 is functionally inactive and can phenocopy SRSF1 loss.^54^ ^55^ SRPK1 phosphorylates serines within Arginine-Serine 1 (RS1) domain to drive nuclear entry, and CLK1 modifies serines within RS/Pro-Ser repeats in RS2 to enable pre-mRNA engagement.^56^ Accordingly, knocking down SRPK1 perturbs SRSF1-dependent AS events.^57^ ^58^ ^59^ We first asked whether altered SRSF1 abundance explains the RI phenotype. To this end, we compared transcript levels of SRSF1, SRPK1 and CLK1 in patient samples with SF mutations and healthy volunteer controls. CLK1 levels were modestly downregulated across all three SF mutant groups (-0.65 to -0.75 L2FC, FDR <0.05) in both datasets,^13^ ^12^ while SRSF1 and SRPK1 levels were unchanged (**Fig. 4A; Fig. S4A**). This prompted us to test whether alterations in SRSF1 function may arise due to changes in phosphorylation status, rather than abundance.

**Figure 4:**
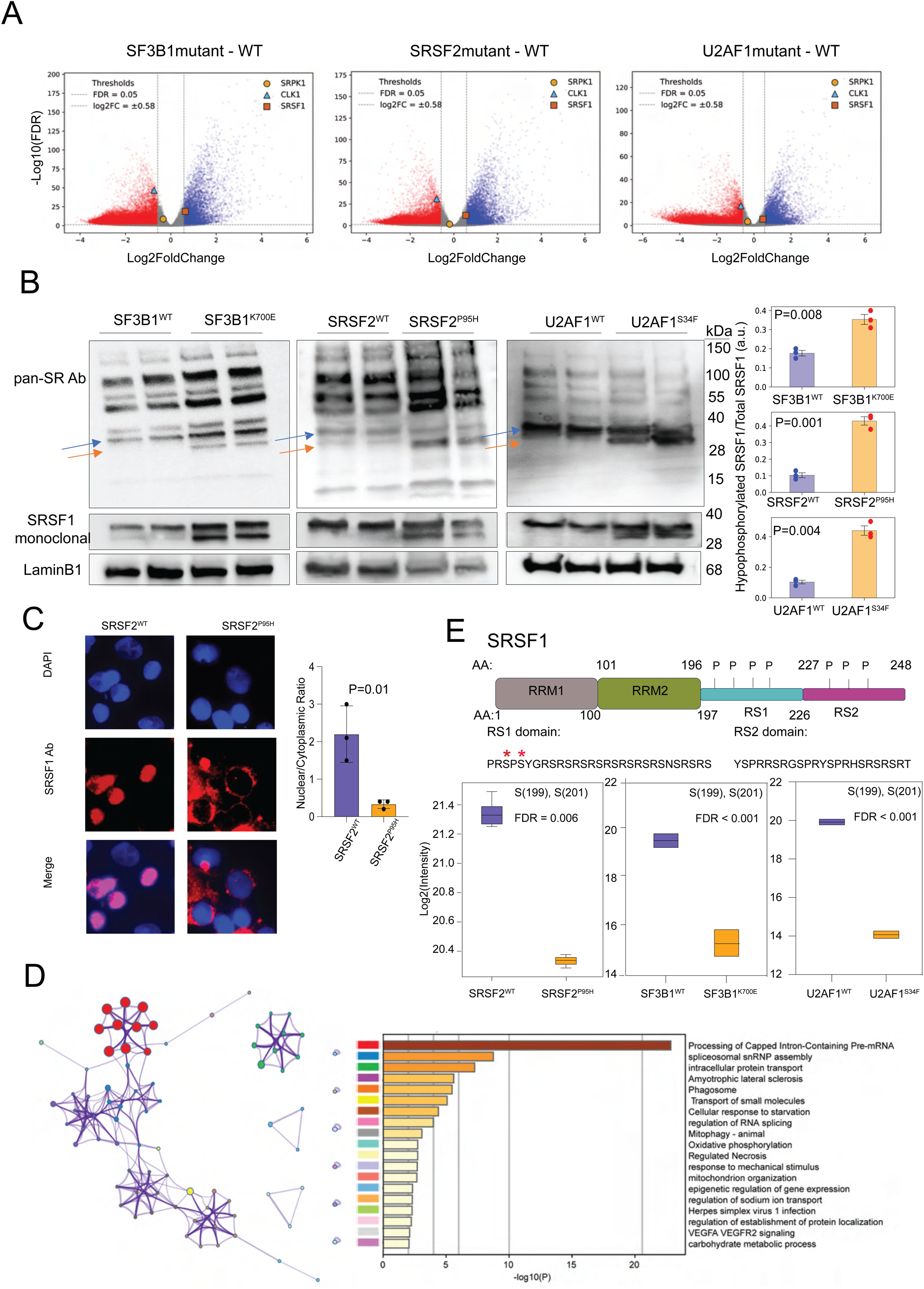
SRSF1 phosphorylation changes in SF-mutant cell lines. (A) Volcano plots illustrating differential gene expression in SF mutant versus WT conditions, based on data from Hershberger et al. Each point represents a gene, with its x-axis value indicating the Log2 fold change (Log2FC) and its y-axis value indicating the -log10 of the false discovery rate (FDR). Highlighted genes, including SRPK1, CLK1, and SRSF1, are annotated in the plots to indicate their positions relative to the thresholds. (B) Western blot analysis shows increased hypophosphorylated SRSF1 (orange arrows) in splicing factor mutants compared to WT, as detected by pan-SR antibody and SRSF1 monoclonal antibody. Total SRSF1 levels remain unchanged. Quantification of intensities by ImageJ confirms significant increases in hypophosphorylated SRSF1 levels in mutants. (C) Immunofluorescence analysis of SRSF1 in SRSF2^P95H^ and SRSF2^WT^ cells. Cells were stained with DAPI (blue) to visualize nuclei and immunostained with anti-SRSF1 antibody (red). Right, quantification of nuclear-to-cytoplasmic SRSF1 signal intensity ratio (n = 3 replicates). Bars represent mean of medians. SRSF2^P95H^ cells show significantly reduced nuclear enrichment of SRSF1 compared to wild-type controls (p <0.001 by two-tailed t-Test). (D) Metascape analysis of mass spectrometry phosphoproteomic data between SRSF2^P95H^ and SRSF2^WT^ conditions. Network Plot (Left) showing clusters of enriched biological processes with nodes representing terms and edges indicating shared genes. Nodes represent protein phosphosites and colors represent the pathways they belong to. Bar Plot (Right): Top pathways ranked by significance (-log10(P)), highlighting RNA splicing, intracellular transport, and mitochondrial organization. (E) Schematic of SRSF1 domains and phosphorylation analysis. The RS1 domain (amino acids 197-226) contains phosphorylation sites S199 and S201. Box-whisker plot shows log2 intensity of S199/S201 phosphorylation in (purple) vs. (orange). Phosphorylation is significantly reduced across all 3 mutant conditions (SRSF2, n=3 replicates each; SF3B1 and U2AF1, n=2 replicates each).

We utilized doxycycline-inducible K562 cell systems, including endogenously edited SF3B1^WT^ and SF3B1^K700E^,^60^ as well as lentiviral overexpression models for SRSF2^WT^ and SRSF2^P95H^, and U2AF1^WT^ and U2AF1^S34F^.^61^ Given that SRSF1 is a highly basic protein, phosphorylation can significantly lower its isoelectric point,^62^ leading to migration shifts in gel electrophoresis. Western blot analysis using SRSF1-specific and pan-phospho-SR antibodies (**Fig. 4B**) showed a significant increase in the hypophosphorylated form of SRSF1 across all three SF mutant states, with no change in total SRSF1 levels. Consistently, immunofluorescence (IF) revealed significant decrease in the nuclear-to-cytoplasmic SRSF1 signal ratio in the mutant conditions (**Fig. 4C**). Thus, SF-mutant protein expression attenuates SRSF1 function via hypophosphorylation and altered nuclear-cytoplasmic distribution.

Having confirmed changes to SRSF1 phosphorylation, we then profiled phosphoproteomes of K562 cell lines expressing either wild-type or mutant splicing factors, with a focus on SRSF2^WT^ vs. SRSF2^P95H^ cell lines for detailed comparison (N = 3, each). Using a library-free directDIA workflow (Methods), we identified 52,619 unique phosphopeptides (46,019 ± 5,659), mapping to 34,890 unique class-I phosphorylation sites (or, P-sites, with localization probability > 0.75) assigned 6005 unique phosphoproteins in the entire experiment, with the protein-FDR controlled below < 0.01 (**Supplementary Excel 2**).^63^ Replicates were highly reproducible (median absolute Pearson correlation was .99 and .96 within the SRSF2^P95H^ and SRSF2^WT^ sample replicates, respectively) (**Fig. S4B**). Both PCA (**Fig. S4C**) and hierarchical clustering (**Fig. S4D**) separated samples by mutation status (SRSF2^P95H^ vs. SRSF2^WT^), indicating mutation status as the key driver for phospho-signaling changes. Metascape^64^ pathway analysis highlighted a significant upregulation of pathways of RNA splicing and processing, such as ‘Processing of Capped Intron-Containing Pre-mRNA’ and ‘Spliceosomal snRNP Assembly’ (**Fig. 4D**). Pathways associated with cellular adaptation to metabolic stress, such as ‘Cellular Response to Starvation,’ ‘Mitophagy,’ and ‘Oxidative Phosphorylation,’ were also noted to be upregulated in SRSF2^P95H^. The extensive RS repeats in SRSF1’s RS2 generate numerous trypsin cleavage sites, limiting accurate LC-MS/MS profiling of this region. MS was however able to effectively capture changes at the N-terminal of the RS1 domain of SRSF1; Ser199 and Ser201 phosphorylation was significantly downregulated in all mutant conditions-SRSF2 ^P95H^, SF3B1^K700E^ and U2AF1^S34F^ without a change in the corresponding total SRSF1 levels (**Fig. 4E, Fig. S4E**). Importantly, these serines are known targets of SRPK1^65^ and AKT,^66^ ^67^ suggesting impaired activity of these upstream kinases. Collectively, these findings reveal that SF-mutant states do not lower SRSF1 abundance but instead shift SRSF1 to a hypophosphorylated, cytoplasm-biased state, accompanied by site-specific loss of SRSF1 phosphorylation consistent with diminished SRPK1/AKT signaling.

### Replicative stress disrupts AMPKα/AKT balance, nuclear speckle organization, and induces SRSF1 hypophosphorylation in SF-mutants

Based on MS evidence of reduced SRSF1 phosphorylation, we examined how these changes may fit with AKT and AMPKα activity. Several canonical AKT substrates^63^ were significantly downregulated, consistent with an overall attenuation of AKT/mTOR signaling (**Fig. 5A; Fig. S5A**). Among these,= was mTOR-Ser2481, a marker of intact mTORC2 activity,^68^ which plays a critical role in phosphorylating AKT1 at Ser473 (**Fig. S5B**). The only upregulated AKT substrate was NOS3-Ser1177 phosphorylation; however, as a target of both AKT^69^ and AMPKα,^70^ this likely reflected AMPKα activation, which is metabolically antagonistic to AKT.

**Figure 5:**
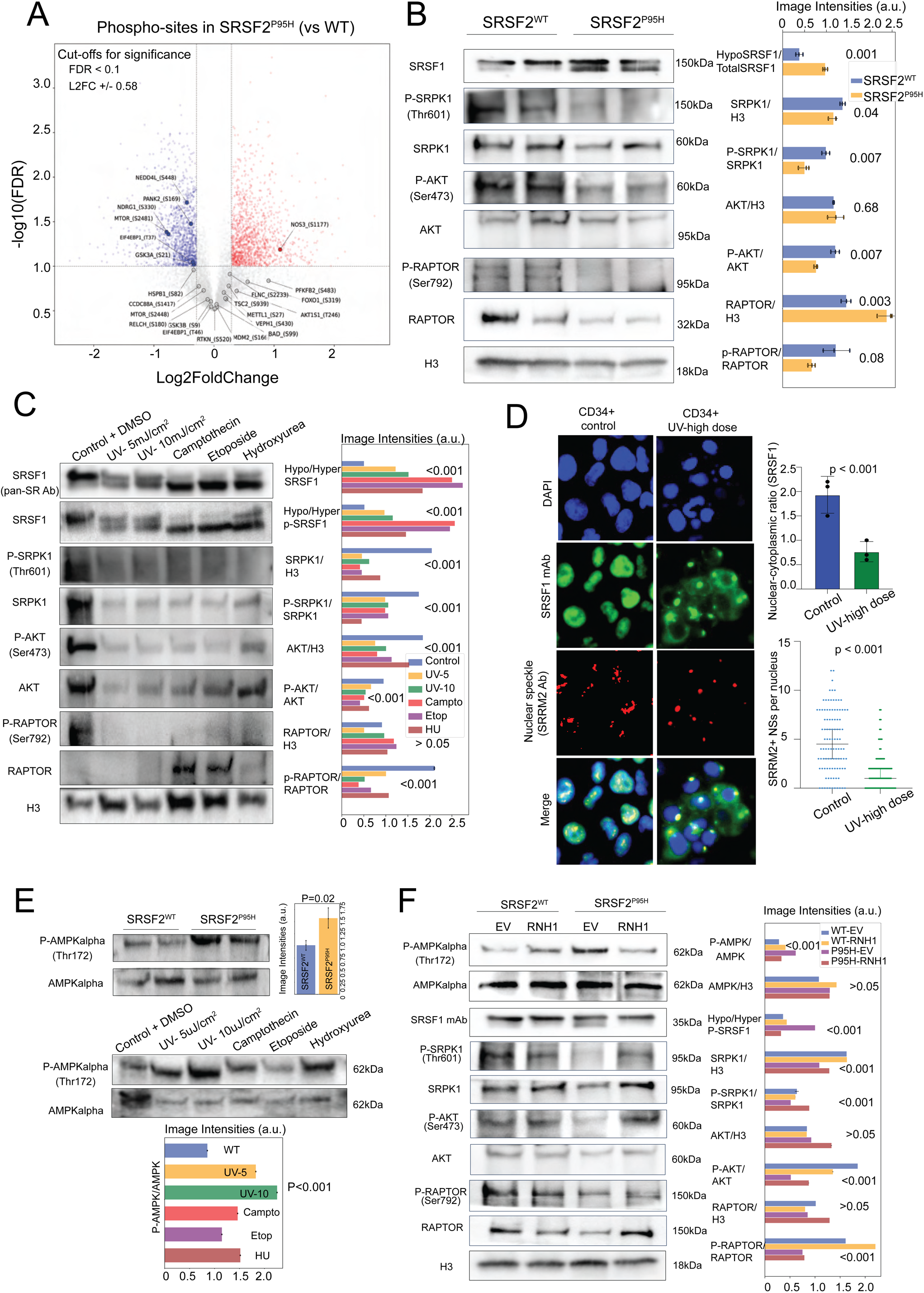
DDR and replicative stress induce SRSF1 phosphorylation changes similar to that observed in SF mutant cell lines. (A) Volcano plot of AKT-target phosphosites in SRSF2^P95H^ vs SRSF2^WT^. Differentially phosphorylated sites are highlighted with significant fold change (L2FC±0.58 and FDR < 0.1). Key AKT-regulated upregulated (red) and downregulated (blue) phosphosites are annotated. (B) Western blot analysis comparing SRSF2^P95H^ vs SRSF2^WT^ cell lines for phosphoprotein levels. Quantification of image intensities for key ratios highlights significant differences, with p-values annotated for statistical comparisons. a.u: arbitrary units. (C) Western blot analysis of phosphoprotein levels under various treatments, including UV irradiation (5 mJ/cm² and 10 mJ/cm²), camptothecin (Campto), etoposide (Etop), and hydroxyurea (HU). Quantification of key protein ratios is displayed as image intensities (a.u.) (mean value of triplicates) with notable differences between treatments compared to the DMSO control. (D) UV-induced relocalization of SRSF1 and loss of nuclear speckles in primary CD34^⁺^cells. Left, representative IF of untreated (control) and UV-high-dose-treated CD34^⁺^ hematopoietic progenitors stained with DAPI (blue), SRSF1 (green), and the speckle marker SRRM2 (red); bottom row shows merged channels. Right, quantitative analyses: (top) nuclear-to-cytoplasmic SRSF1 signal ratio (mean of medians, n = 3 fields per condition); (bottom) number of SRRM2-positive speckles per nucleus (each dot = one nucleus, mean ± SEM). UV treatment significantly reduces nuclear SRSF1 enrichment and nuclear speckle (NS) count (two-tailed t-test and Mann-Whitney U test, p < 0.001 for both metrics). (E) Differential phosphorylation of AMPKα at Thr172 in SRSF2^P95H^ vs SRSF2^WT^ cells (Top panel), and CD34+ control and stress conditions (Bottom panel; as in Fig. 4C). Quantification of P-AMPK/AMPK ratios shows increased phosphorylation in mutant conditions compared to WT. Bar plots represent mean intensities ± SD from three replicates. Statistical significance indicated. (F) RNase H1 expression partially rescues phosphorylation defects in SRSF2^P95H^ mutant cells. Western blot analysis shows phosphorylation levels of key signaling proteins and their corresponding total protein levels in SRSF2^WT^ and SRSF2^P95H^ cells transduced with either empty vector (EV) or RNase H1. Quantification of phosphorylation and protein ratios presented as bar plots with mean ± SEM (triplicates).

AKT1 can phosphorylate SRSF1 either directly within its RS1 domain^66^ ^67^ ^71^ or indirectly by activating SRPK1, which in turn phosphorylates SRSF1.^72^ Immunoblotting confirmed reduced activation of AKT1 (decreased AKT1-Ser473)^73^ ^74^ and diminished phospho-SRPK1 levels across the three SF mutant conditions (**Fig. 5B; Fig. S5C**). Reduced phosphorylation at both mTOR-Ser2481 (mTORC2 component) and Raptor-Ser792 (mTORC1 component)^75^ ^76^ (**Fig. 5B**) was also seen, suggesting a global downregulation of mTOR signaling, affecting both mTOR complexes.

To determine the mechanisms driving these changes to the kinases, we explored potential upstream pathways commonly shared across the three SF mutant conditions. In this context, multiple studies have reported elevated R-loop induced replicative stress and DNA Damage Response (DDR) across *SF3B1*, *U2AF1*, and *SRSF2* mutations.^77^ ^78^ Using S9.6 antibody slot blot, we first confirmed R-loop in all three SF mutations (**Fig. S5D**). Next, we determined if replicative stress with various DDR-inducing agents could similarly induce SRSF1 hypophosphorylation. Because transformed cell lines such as K562 undergo rapid apoptosis in response to DDR induction, we used primary CD34+ derived hematopoietic progenitors. Cells were treated with various DNA-damaging agents, including UV radiation (single-stranded DNA damage),^79^ camptothecin and etoposide (topoisomerase inhibition),^80^ and hydroxyurea (replication fork stalling),^81^ all of which reliably activate DDR. All four DDR triggers reduced the phosphorylation of SRSF1, Raptor at Ser792 and AKT1 at Ser473 (**Fig. 5C**). Total levels of both phospho-SRPK1 and SRPK1 were also reduced. IF showed a striking redistribution of SRSF1 from into the cytoplasm (**Fig. 5D**), accompanied by a marked reduction in the number of nuclear speckles (NSs), themselves. Because NSs store and release phosphorylated SR proteins, we interpret this to be consistent with loss of the phosphorylated SR-protein pool and impaired co-transcriptional splicing. Together, DDR activation closely mirrored changes to the SRSF1 and its regulatory kinases in SF-mutant cell lines.

We next asked what links DDR to mTOR/AKT axis and SRSF1 hypophosphorylation? DDR activates several protein kinases, including AMPKα,^82^ ^83^ ^84^ a major metabolic stress sensor that promotes catabolic signaling and suppresses anabolic pathways. AMPKα and AKT are functionally antagonistic: AMPKα activation inhibits AKT phosphorylation at residues Ser473 and Thr308,^85^ thereby suppressing AKT activity. Conversely, active AKT is associated with reduced AMPKα phosphorylation at its major regulatory site, Thr172, on the α-catalytic subunit.^86^ ^87^ Consistent with the patterns expected in DDR-induced metabolic stress, we observed increased AMPKα-Thr172 phosphorylation in both DDR-activated CD34+ cells and SF mutant cells (**Fig. 5E**). These findings suggest that DDR-induced metabolic stress disrupts the relative balance of AMPKα and AKT activity by AMPKα-Thr172 upregulation and AKT-Ser473 downregulation. This imbalance in turn inhibits the AKT-SRPK1-SRSF1 axis, leading to hypophosphorylation of SRSF1.

We next evaluated whether pharmacologic DDR activation reproduces the SF-mutant RI signatures. Because short-term induction changes may not yet be reflected in total steady-state RNA, we instead sequenced newly synthesized RNA (4-hour 4-thiouridine exposure; Methods) after acute induction of DDR in CD34+-derived cells and compared these profiles to the physiological baseline. Across 4,918 introns, DNA damage/stress treatments captured a broad share of the SF3B1-mutant RI program (**Fig. S5D**). At |dIR| ≥ 0.02, etoposide showed the largest sign-agnostic overlap with SF (4,635 introns), followed by hydroxyurea (4,051), camptothecin (3,917), UV_high dose (3,549) and UV_low dose (2,902) (**Fig. S5E**). Directionally, hydroxyurea most closely mirrored SF alteration state, with 52.4% of shared events changing in the same direction (95% CI ∼50.9-54.0), while camptothecin and UV treatments were intermediate (∼48-50%) and etoposide lower (46.2%) (**Fig. S5F**). These rankings were stable across different dIR thresholds (|dIR| ≥ 0.01-0.10). A large, shared core emerged: 2,178 introns were altered by all five treatments and SF. Within this core, 46.4% showed majority concordance with SF (≥ 3/5 treatments aligned; binomial p=0.34 vs null: 0.5), and 2.5% were strictly concordant across all five (p=0.12 vs null: 3.125%), indicating that while the same introns are stress-sensitive, the direction of regulation is context-dependent. For instance, upon intersecting the UV_high dose, camptothecin, and SF subsets (3,106 introns), 24.2% moved in the same direction (**Fig. S5G**), indistinguishable from the 25% expected by chance (p=0.34). Collectively, these data position SRSF1 hypophosphorylation as a shared entry point into the RI program across DDR conditions, but not its sole determinant. Although DDR treatments and the SF-mutant state target many of the same introns, the direction of change often diverges, both across DDR conditions and relative to the SF-mutant state. This suggests the presence of DDR-specific regulatory layers, including perturbation of kinase and RBP families, that remain unaffected in SF mutants but still shape splicing outcomes.

Finally, we determined if alleviating replication stress and downstream DDR could potentially restore the AMPKα-AKT balance. We reversed R-loop induced DDR by overexpression of RNAseH1^88^ using a doxycycline-inducible lentiviral system in SF mutant cells. RNaseH1-induced reduction of R-loops (**Fig. S5H**) was accompanied by decreased AMPKα-Thr172 phosphorylation, increased AKT-Ser473 phosphorylation, and increased SRSF1 phosphorylation levels (**Fig. 5F, Fig. S5I**). Collectively, these findings support a model in which DDR-driven metabolic stress suppresses mTOR/AKT, impairs the AKT-SRPK1-SRSF1 axis, reorganizes nuclear speckles, and ultimately rewires splice-site choice.

### PI3K/AKT pathway and SRSF1 phosphorylation are altered in SF-mutant patient samples

We next sought to assess AMPKα activation and AKT inhibition in primary AML/MDS samples with SF mutations. Bone marrow samples of patients with relatively high clonal component (VAFs >= 20%) were chosen. To reduce confounding variables, we also prioritized those without co-existing level I variants in non-SF genes. Because each cryopreserved BMMNC vial yields 25-50,000 cells, insufficient for immunoblotting low-abundance phospho targets, we broadened sampling to committed progenitors. MDS is a stem cell disorder with driver mutations across various stages of myeloid differentiation including such committed progenitors.^89^ We therefore also included CD34^low/-^ CD41^+^ (myeloid cells committed to megakaryocyte lineage)^90^ and CD34^low/-^CD15^+^ (myeloid cells committed to granulocyte lineage) populations.^91^ This approach provided up to 0.25-0.5 million total cells (**Fig. 6A**), sufficient to evaluate the phosphorylation status of multiple proteins. Western blot analyses revealed increased AMPKα-Thr172 phosphorylation, reduced Raptor-Ser792 and AKT-Ser473 phosphorylation, and a marked downregulation of both phosphorylated and total SRPK1 levels across multiple patient samples (**Fig. 6B**). Significant hypophosphorylation of SRSF1 and a pronounced reduction in total SRSF1 protein levels were also seen, particularly in the two SRSF2-mutant cases. Peripheral blood mononuclear cells from a patient with circulating SF3B1-mutant cells similarly showed SRSF1 hypophosphorylation (**Fig. 6C**), indicating that these changes extend to more differentiated compartments. We next performed transcriptome analysis (RNA-seq-derived fold changes) in *SF3B1*, *U2AF1*, and *SRSF2* mutants (compared to the healthy volunteer controls) ^13^ using Ingenuity Pathway Analysis (IPA).^92^ The top canonical pathways identified as significantly activated or inhibited in the SF mutations are illustrated in **Fig. S6A**. The PI3K/AKT signaling pathway was consistently predicted to be inhibited across all three mutant conditions, as evidenced by negative z-scores in IPA (**Fig. S6A, Fig. S6B**).

**Figure 6.**
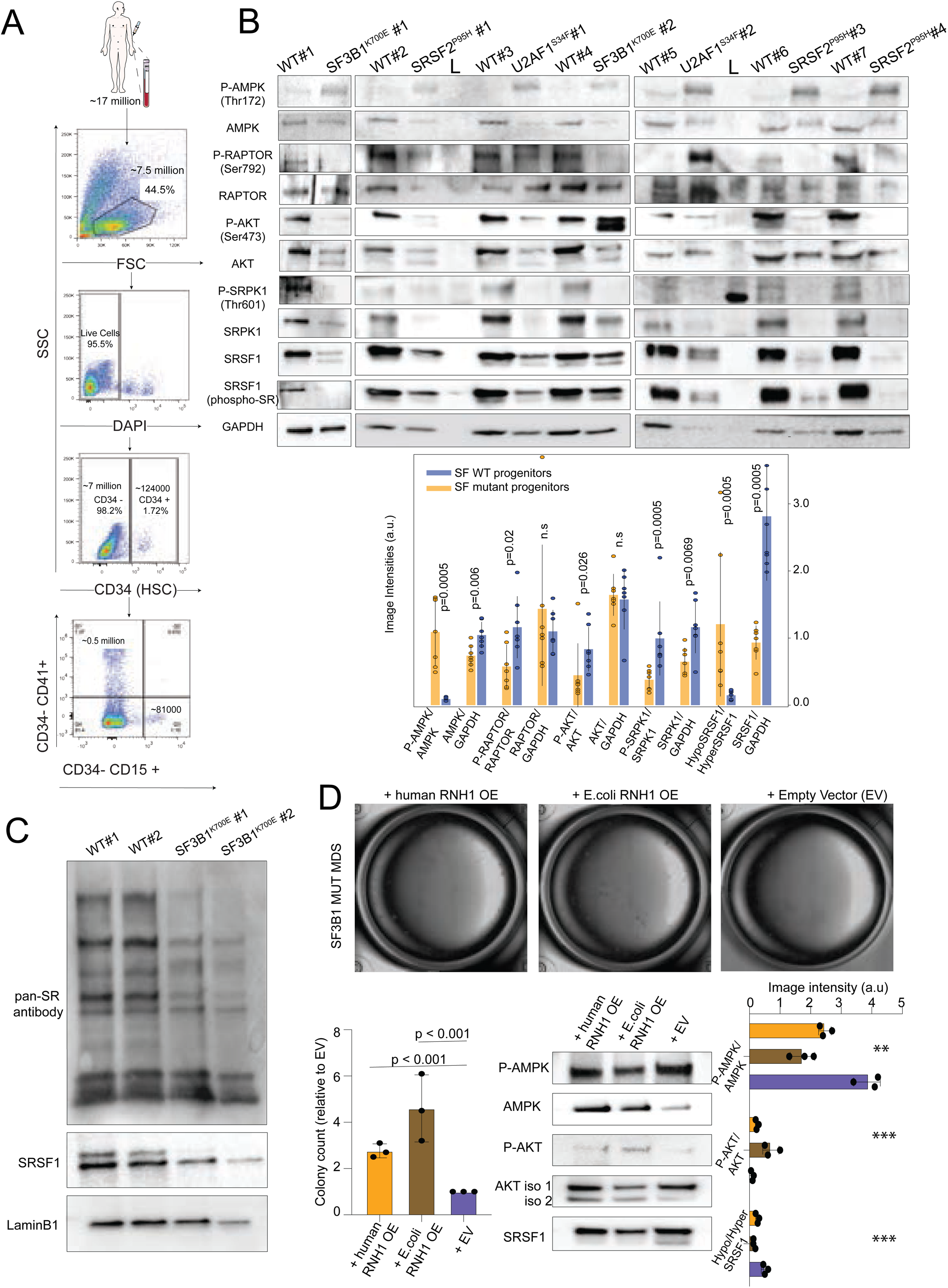
PI3K/AKT pathway downregulation and SRSF1 protein and phosphoprotein levels in patient-derived progenitor cells. (A) Flow cytometric gating strategy for cell subset isolation. CD34+ HSCs accounted for ∼1.72% (∼124,000 cells), with ∼7 million CD34− cells (98.2%). Further gating identified CD34^−^CD41+ and CD34^−^CD15+ committed progenitors. (B) Western blot analysis of phosphorylation and total protein levels in SF WT and SF mutant progenitor cells. GAPDH used as a loading control. Quantified image intensities for phosphorylated and total protein levels are shown in the bar graph, comparing SF WT (blue) and SF mutant (orange) progenitors. Data are presented as mean ± SEM, with p-values calculated for statistical significance. Abbreviations used: n.s.-not significant, a.u-arbitrary units, L-protein ladder. (C) Western blot analysis of extracts from WT and K700E mutant peripheral blood mononuclear cell samples. Top: pan-SR antibody detects the phosphorylation status of SR proteins. Middle: SRSF1-specific antibody shows total SRSF1 levels. Bottom: LaminB1 serves as a nuclear loading control. The reduced signal in the SF3B1^K700E^ lanes for the pan-SR antibody indicates hypophosphorylation of SRSF1. (D) Representative colony-forming unit (CFU) assays of pooled CD34^⁺^ cells from SF3B1^K666Q^ MDS patient samples transduced with human RNase H1 (human RNH1 over expression (OE)), *E*. coli RNase H1 (E. coli RNH1 OE) or empty vector (EV). Plates were scored after 14 days. Lower left: Quantification of colony counts normalized to EV (mean ± SEM, n = 3 independent observations; p < 0.001 by one-way ANOVA with Tukey’s post hoc test). Lower right: Western immunoblots for P-AMPKα (Thr172), total AMPKα, P-AKT (Ser473), AKT and SRSF1 in the same transduced samples along with densitometric analysis of P-AMPKα/AMPKα, P-AKT/AKT and Hypo/Hyper SRSF1 ratios (mean ± SD, n = 3). \***\***Statistics by one-way ANOVA.

We next tested whether relieving R-loop-driven DDR restores AKT-SRSF1 signaling and improves clonal fitness in SF-mutant MDS cells. We overexpressed (OE) human or *E*. coli RNase H1 (RNH1 OE) in pooled CD34⁺ progenitors from a SF3B1^K666Q^ patient sample and performed colony formation assays alongside immunoblot analysis of the phospho targets. Compared to empty vector controls, human RNH1 OE and *E*. coli RNH1 OE increased colony counts by 3x and 4.5x, respectively (both p < 0.001) (**Fig. 6D**), indicating a robust rescue of proliferative capacity, consistent with previous studies.^77^ ^78^ Furthermore, immunoblots of expanded cells revealed that RNH1 OE partially reversed AMPKα-Thr172 hyperphosphorylation, restored AKT-Ser473 phosphorylation to near-control levels, and rescued SRSF1 phosphorylation (**Fig. 6D**). Sanger sequencing confirmed retention of the mutation alleles in colonies across conditions (**Fig. S6C**), indicating that the RNH2 OE rescue reflected mutant clone outgrowth, and not contamination from expansion of normal/WT progenitors. Finally, we determined if resolving R-loops would shift the splicing pattern (as suggested by our results that showed strong correlation of SF-mutant splicing patterns with SRSF1 knockdown and anticorrelation with hnRNPA0 depletion, **Fig. 3C**). We therefore performed RNA-seq on SF3B1^K666Q^ progenitors expressing EV, human RNH1 OE, or *E*. coli RNH1 OE. For each sample, we quantified PSI for RI, SE, A3, and A5 events, then computed ΔPSI (OE - EV). We clustered these two ΔPSI profiles alongside the various shRNA K562 KD conditions (**Fig. S6D**), by Ward linkage on Euclidean distances. Across all AS classes, both human and E. coli RNH1 ΔPSI signatures co-clustered with hnRNPA0 and hnRNPAB knockdowns. Together, these data demonstrate that R-loop resolution mitigates DDR-induced suppression of the AKT-SRPK1-SRSF1 axis, restores clonal outgrowth in SF-mutant MDS progenitors, and changes splicing pattern closer to baseline.

## Discussion

In this study, we show that DDR-induced changes in phosphorylation of RBPs such as SRSF1 represent a key mechanism regulating splicing outcomes in SF-mutant states. While the handful of splicing factors recurrently mutated in cancer each exhibit largely distinct splicing patterns, reflecting the different stages of catalysis in which they function, a notable proportion of altered AS events, particularly within the RI program, are shared across all SF-mutant states. The basis for this overlap was previously unclear. Our results link it to an indirect consequence of replicative stress, which is consistently observed in cells harboring *SF3B1*, *U2AF1*, or *SRSF2* mutations. Thus, the AS landscape in SF-mutant contexts is shaped by both mutant-specific cis effects and shared indirect influences, including those mediated through DDR.

We focused on RI alterations to pinpoint RBPs and kinase pathways driving AS defects, as RI was the most consistently shared across SF mutation categories. Although RBP dysregulation could be expected to broadly affect all AS types, SRSF1 knockdown most strongly mirrored SF mutation-associated changes in RI. This suggests that RI regulation may rely on relatively simple networks dominated by RBPs such as SRSF1 and hnRNPs, whereas other AS modes likely involve more complex, multilayered regulation. RI, once viewed as mere “splicing failure,” is now recognized as a regulated mode of gene expression.^93^ ^94^ Retained introns are typically nuclear and can serve as a reservoir for rapid splicing and export when needed.^44^ While prior studies have suggested a predominant reduction in RI in SF-mutant states,^13^ ^95^ our analysis revealed both gains and losses, with directionality always concordant across *SF3B1*, *U2AF1*, and *SRSF2* mutants-pointing to a shared mechanism. Comparative analysis with public datasets implicated loss of SRSF1 and gain of hnRNPA0/AB function. Using phosphoproteomics and RNA-seq, we connect reduced SRSF1 activity to DDR-induced imbalance of AMPKα/AKT signaling, characterized by AMPKα-Thr172 activation and AKT-Ser473 inhibition (**Fig. 7**), a pattern consistent with cellular adaptation to metabolic stress. Various metabolic stress responses could potentially induce a similar AMPKα-AKT signaling pattern.^22^ Alleviating replication stress by overexpressing RNaseH1 reversed AMPKα/AKT signaling defects and restored SRSF1 phosphorylation, indicating that DDR is a primary driver of SRSF1 hypophosphorylation. Concomitantly, we observed reduced SRSF1 phosphorylation across various DNA damage-inducing agents, underscoring the link between DDR and RBP activity, including that of SRSF1. DDR in SF-mutant states can be reversed by over-expression of RNAseH which resolves R-loops. We show that reversing DDR by over-expressing RNAseH can also reverse changes to SRSF1 phosphorylation and RI in the transcriptome, providing further evidence to the link between DDR and RI. Studies in patient samples mirrored many aspects of this connection. Building on previous studies that describe increased R-loops and replicative stress as hallmarks of *SF3B1*, *U2AF1*, and *SRSF2* mutations,^96^ ^97^ ^98^ ^99^ we investigated DNA damage as a major metabolic stressor influencing SRSF1 function in SF mutant cells. While the link between DDR and SRSF1 through AMPKα/AKT is what we focused on, effects of DDR or other cellular stress is likely complex with many layers of regulation. DDR may not be the only form of stress in SF mutant states and many others such as unfolded protein response maybe operant in such scenarios. Splicing is also intricately linked to transcription, which is affected by DDR and cellular stress providing yet another layer of regulation-our group has shown previously how SF mutations reduce Pol II elongation rate,^100^ which may affect splice site choice. Unbiased systems-based approaches which incorporate more comprehensive profiling of post-translational changes to RBPs and long read sequencing for RI will dissect this complex web of regulation.

**Figure 7.**
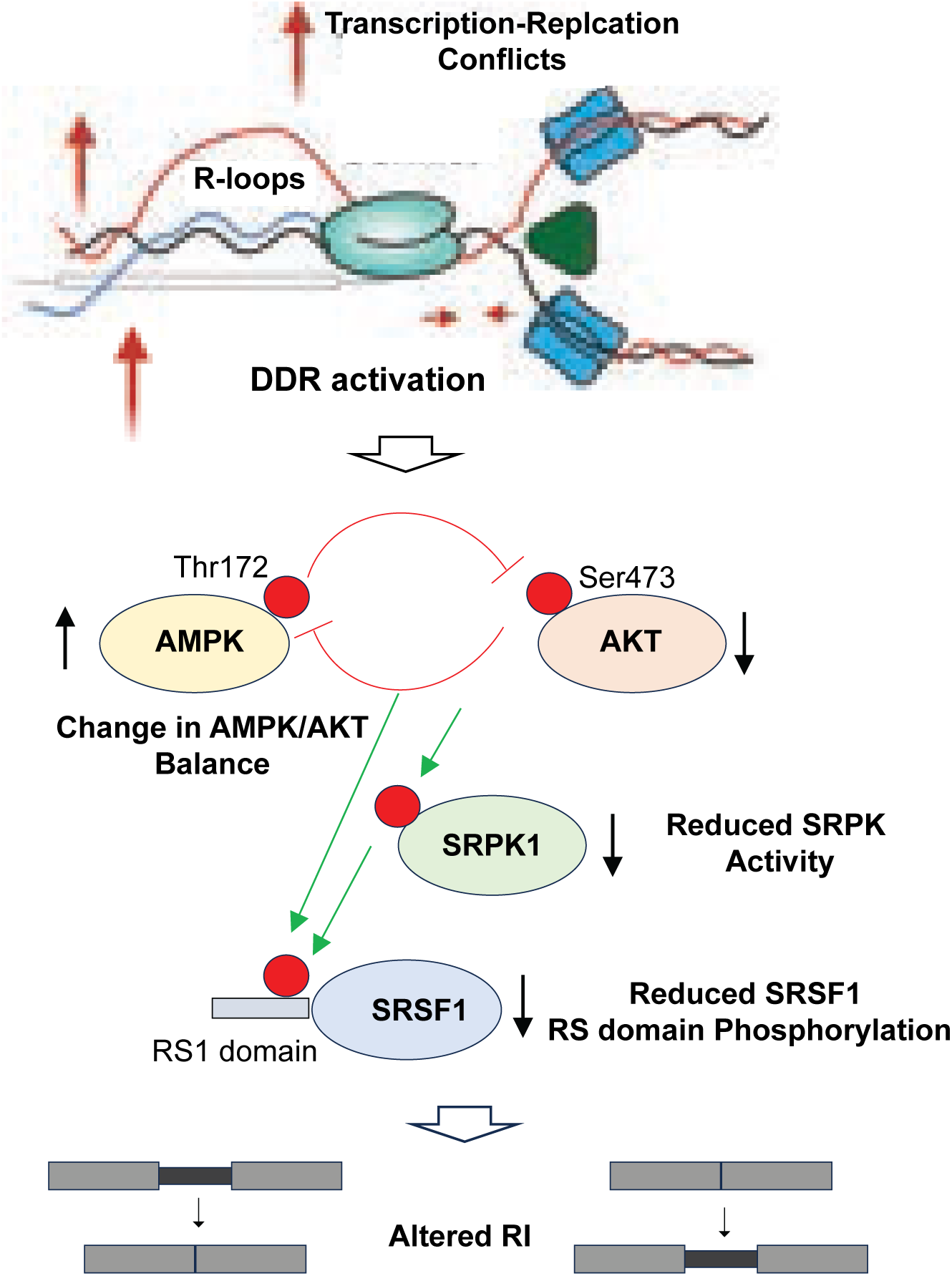
Schematic representation of the signaling cascade linking transcription-replication conflicts to DDR activation. Increased R-loops and DDR stress lead to phosphorylation changes at key sites, including AMPK (Thr172) and AKT (Ser473). These alterations modulate downstream kinases such as SRPK1, affecting SRSF1 phosphorylation at the RS1 domain. Green arrows indicate activation.

Our observation of SRSF1 hypophosphorylation by DDR contrasts with a report^101^ of increased phosphorylation after UV or high-dose etoposide. This discrepancy likely reflects experimental differences (dose, cell type, or duration of DDR); it I also possible that DDR imparts complex changes with both hyper- and hypo-phosphorylation of specific residues, producing a net functional disruption. Nevertheless, a consistent reduction of phosphorylation across multiple DDR-inducing strategies, together with mass spectrometry evidence of RS1 domain dephosphorylation, supports a central role for SRSF1. Interestingly, patient transcriptome revealed a modest but significant downregulation of CLK1 expression-CLK1 is known to regulate RI^102^ and hence could be another contributor to the observed changes in SRSF1 phosphorylation and RI. Finally, our results showed a reversal of RI upon reversal of replication stress by multiple approaches. Many of the implicated signaling pathway proteins, such as AMPKα^103^ and AKT^104^ are druggable, and open the avenue for pharmacologic intervention to reverse the phenotype of MDS.

In summary, our study shows that widespread changes to alternative splicing in SF-mutant states is not limited to direct cis effects of the neomorphic mutant protein. Indirect effects from cellular stress or transcriptional speed affect the final transcriptional output. Since correcting cellular stresses such as those from DDR can restore the aberrant transcriptome, it opens avenues for therapeutic interventions focusing on these signaling pathways.

### Limitations of the study

We recognize that RBP effect on splicing is likely complex and operates through interconnected networks,^105^ rather than being limited to a single factor, such as SRSF1, which was our focus here. Our study also did not address how stressors other than DDR may impact AS. RI as determined by short read sequencing is limited by multimapping of reads in repeat regions and poor coverage of very long introns. Mass Spectrometry data is limited by poor capture of RS repeats after trypsin digests, which prevented full profiling of RS domains, leaving gaps in broader understanding of changes to SRSF1 phosphorylation dynamics.

## Methods

### Alternative splicing analysis

Transcript abundance estimation was performed using Salmon v1.10.2 (Ref) with the pseudo-alignment method, employing a Salmon index constructed from Gencode release v46. For the analysis of alternative splicing (AS) events, SUPPA2 v2.4 was used. First, AS events were generated using the Gencode v46 GTF file with the generateEvents option in SUPPA2, creating input ioe files for five types of AS events: mutually exclusive exons (MXE), skipped exons (SE), alternative 5’ splice sites (A5), alternative 3’ splice sites (A3), and retained introns (RI). The generated ioe files and transcript abundance files in transcripts per million (TPM) units from Salmon were used to calculate the relative inclusion level (PSI) of each AS event per sample. Differential splicing analysis was conducted across all AS types in a single integrative approach between the various splicing factor mutant conditions and the wild-type control using the diffSplice function in SUPPA2. The analysis was executed with the following parameters: --method classical, --area 1000, -me, -gc, and --tpm-threshold 5 -al 0.05. Events were considered significant if they exhibited an absolute PSI difference greater than |0.1| and an adjusted p-value of less than 0.05.

To visualize the distribution of AS events across different splicing factor mutant conditions, a custom Python script was used to parse the SUPPA2 differential splicing output. Events were stratified by AS type-SE, MXE, A5, A3, and RI- and categorized based on statistical significance. Events were defined as significant if they exhibited an absolute ΔPSI ≥ 0.1 and an adjusted p-value ≤ 0.05. Among these**, ‘**shared & significant’ events were those meeting significance criteria across all seven SF mutant conditions, while ‘significant but not shared’ events were those found to be insignificant in at least one condition. Events that did not meet the significance threshold were classified as ‘non-significant’. The total, shared, unique, and non-significant events were quantified per condition and AS type, and their proportions visualized using a stacked bar plot (**Fig. 1C & Fig. S1D**) to highlight the relative contributions of splicing changes across conditions.

### Slope Calculation and Baseline Stratification

Unspliced intronic reads were quantified in a strand-aware manner using ‘featureCounts’ (paired-end aware; stranded mode). Introns were taken from the IRFinder “clean” intron set (regions not overlapping annotated exons/UTRs). Each clean intron was subdivided into 10 equal-length bins (“deciles”) and counted per library and condition. Because replicate numbers differed between conditions, we summed counts, per decile, across all libraries within a condition and then normalized each decile by that condition’s total intronic reads to obtain relative intronic coverage per decile (averaging per-library first and then normalizing gave concordant results).

To focus on ends of introns we retained deciles 1-2 and 9-10 only. Using the strand annotation, deciles 1-2 correspond to the 5′ end and 9-10 to the 3′ end on the “+” strand, and the mapping is reversed on the “−” strand. We required ≥500 aggregate counts per decile in both conditions to reduce noise. For each intron and condition we computed:

c5 = mean normalized coverage of the two 5′ deciles, c3 = mean normalized coverage of the two 3′ deciles.

The primary summary of the 5′→3′ gradient was the log-ratio slope:

S_log_ = log_2_(c3 + ε/c5 + ε), with ε=10^−9^ to stabilize very small values. By construction, S_log_ < 0indicates 5′-biased coverage (c5>c3) and S_log_ > 0 indicates 3′-biased coverage (c3>c5).

Log-ratio slope was chosen over classic difference (c3 - c5) because it quantifies 3′↔5′ imbalance on a fold-change, symmetric and scale-invariant basis, is stable at low coverage, reduces heteroscedasticity for cleaner Δ∼WT residualization. For each intron we formed the change in slope between conditions, ΔS □= □S_mut_ − S_WT_. Because ΔS is arithmetically coupled to S_WT_ (features that start far from zero tend to move toward zero even by chance), we residualized ΔS against S_WT_ using ordinary least squares across introns:

ΔS_resid_ = ΔS − (*a*⋅S_WT_+*b*), fitting ΔS ∼ (*a*⋅S_WT_+*b*) across introns. Residualization assumed a linear Δ∼WT trend (empirically satisfied in our data). Intuitively, ΔS_resid_ is “the change beyond what is expected from the WT starting value.” All hypothesis tests below used ΔS_resid_.

WT introns can be either 5′-biased (negative S_WT_) or 3′-biased (positive S_WT_). A flattening means “moving toward zero” in both cases, but that’s a negative residual change for one class and positive for the other. To make interpretation uniform, we sign-aligned residuals by the WT bias:

ΔS_aligned_ □= □ΔS_resid_ × sign(S_WT_)

After this step, ΔS_aligned_ < 0 always means “flatter-than-expected,” ΔS_aligned_ = 0 means no change, and ΔS_aligned_ > 0 means steeper-than-expected, regardless of whether the intron was 5′- or 3′-biased in WT. For instance, slowing co-transcriptional splicing tends to reduce the 5′-3′ contrast by increasing coverage at the previously depleted end (5′ if WT was 3′-biased; 3′ if WT was 5′-biased). The log-ratio log_2_(c3/c5) therefore moves toward 0; sign-alignment maps that flattening to the negative direction for every intron.

Introns were analyzed separately by IR class (LoIR vs GoIR, from the IRFinder). To examine length effects, WT intron lengths were split into tertiles; for the summary figure we collapsed short + mid versus long using the 66.7^th^ percentile of WT length as the long-bin boundary (the boundary is reported from the data). For each IR class and length bin we tested whether the median ΔS_aligned_ was < 0 (flattening) using the Wilcoxon signed-rank test (one-sided). In addition to length, we stratified introns by GC content, predicted fold-energy (FE), 5′ splice-site score (SS5), and 3′ splice-site score (SS3). Attribute values were taken from the WT side (per intron; identical across deciles) and tertile cut-points were computed once across all introns pooled over LoIR+GoIR to ensure common bins. For each attribute we generated All, low, mid, and high panels and, within LoIR and GoIR separately, tested whether the median ΔS_aligned_ was < 0 using a one-sided Wilcoxon signed-rank test (same trimming/plotting conventions as for length; statistics used all values). Reported cut-points are taken from the data. For figures, we plotted histograms with KDE overlays of ΔS_aligned_ for ‘All’, ‘short+mid’, and ‘long’ introns; x-axes were trimmed to the 1st-99th percentiles for display only (tests used all values). We also report overall medians and p-values per group in the text/legends.

To identify independent predictors of the residual, sign-aligned change in slope (ΔS_aligned_), we fit two families of models with heteroscedasticity-robust (HC3) standard errors (SEs). For all models, continuous predictors were standardized (z-scores; mean = 0, SD = 1). We modeled log10 intron length rather than raw length.

1. Linear model (OLS). We regressed the continuous outcome on the standardized predictors and IR class: ΔS_aligned_ ∼ z(log_10_Length) + z(GC) + z(FE) + z(SS5) + z(SS3) + IR_group. Primary inference used HC3-robust SEs and two-sided Wald tests. We also examined IR_group ×predictor interactions using the analogous interaction formula. To quantify unique contribution, we computed drop-one partial R^2^ from nested (classical) OLS fits, and we assessed multicollinearity with variance-inflation factors. Within each table, p-values were adjusted by Benjamini-Hochberg FDR.
2. Logistic model (binomial GLM). For a complementary binary endpoint indicating flattening, y=1{ΔS_aligned_ < 0}.

We fit the same main-effects specification (and, secondarily, IR_group×predictor interactions). Results are reported as Odds Ratios with 95% CIs derived from HC3-robust SEs; discrimination was summarized by AUC. Where indicated, potential non-linearities in z(log_10_Length) and z(SS3) were modeled with natural cubic splines (df = 4), while other covariates entered linearly; predictions were restricted to the observed covariate range. IR_group was treated as a categorical factor (reference level as shown in model tables).

To visualize functional dependence without assuming a parametric form, we plotted LOWESS smoothed curves of ΔS_aligned_ versus z(log_10_Length) separately for LoIR and GoIR, using a smoothing fraction of ∼0.3. The horizontal dashed line marks zero (no change). For annotation, selected z-values were back-transformed to nucleotide units using the dataset’s mean and standard deviation of log_10_length.

### Dimensionality Reduction of Intron Retention Profiles

To assess global patterns of RI, we performed dimensionality reduction on SUPPA2-derived percent spliced-in (PSI) matrices derived from SF-mutant and WT samples,^13^ and hematopoietic lineage-enriched^39^ RNA-seq datasets. PSI matrices were first merged based on common introns and filtered to retain introns with ≥80% completeness across samples. Remaining missing values were imputed using row-wise medians. For PCA, the filtered PSI matrix was transposed (samples as rows) and decomposed using the PCA implementation from scikit-learn (n_components = 2), and principal components were plotted to visualize sample clustering. t-SNE was performed on the RI feature matrix after standard preprocessing. For visualization we first centered and scaled each RI feature across samples and computed principal components; the top 30 PCs were used as input. t-SNE (Barnes-Hut) was run with Euclidean distances, init=’pca’, perplexity = 30 (sensitivity checked at 30-50), learning_rate = max(200, N/12), early_exaggeration = 12, and n_iter = 2,000 with a fixed random seed to ensure reproducibility. Results were stable across seeds and modest parameter changes; the embedding shown was generated with the parameters above.

### Hierarchical clustering of patient samples and ENCODE shRNA knockdown data

The GTF-based SUPPA2 dataset was filtered to include only high-confidence RI events as defined by IRFinder annotations, resulting in a curated set of 4,161 events with robust coverage and minimal overlap with exonic or non-intronic features. To focus on biologically meaningful patterns, we excluded rows with >50% binary inclusion values (0 or 1) or with consistently low inclusion levels (<0.05 across samples). Invalid entries such as NaNs and infinite values were imputed with zeros to maintain computational compatibility during clustering.

For category-based clustering (**Fig. S1F**), patient samples were grouped into predefined categories (e.g., SF3B1 without EM, SF3B1 + EM), and median PSI values were calculated across biological replicates within each group. A distinct color palette was used to distinguish control and mutant categories. Hierarchical clustering was performed using Euclidean distance and complete linkage, chosen to emphasize maximal dissimilarity between categories and effectively separate groups based on the most divergent intron retention patterns. Both rows (RI events) and columns (sample categories) were clustered, and a Spectral colormap was applied to visualize retention gradients across splicing factor mutations.

For hierarchical clustering of ENCODE shRNA knockdown data, we analyzed 143 RBP knockdown conditions in K562 cells, each represented by three biological replicates. Clustering focused on RI events that were consistently altered across three mutation categories, including 2,302 “LoIR” events (delta RI ≥ +0.1) and 1,216 “GoIR” events (delta RI ≤ -0.1), as identified in **Fig. 2A**. For each shRNA knockdown, delta RI values were computed relative to scrambled controls (n = 5) and extracted for the selected RI events. To minimize the impact of outlier magnitudes and emphasize directional similarity, delta RI values were transformed to fixed +0.1 (LoIR) or -0.1 (GoIR). This analysis was exploratory and p-values/FDRs from replicate models were not used to weight distances. Pairwise distances were calculated using the Euclidean metric via SciPy’s ‘pdist’ function, and hierarchical clustering was performed using Ward’s linkage method (to minimize within-cluster variance and well-suited to high-dimensional binary-style data). Clustering results were visualized as dendrograms alongside heatmaps. In Figures **3A**, **S3B**, **S3C** and **S3D**, fixed +0.1/-0.1 values were used to highlight consistent directional effects, while in **Fig. 3B**, actual dRI values were retained to highlight the magnitude of change. A diverging color palette was used in **Fig. 3C** to depict the distribution of dRI values, and the vertical dendrogram reflected the hierarchical relationships among KD conditions.

### Quantification of RI events using IRFinder

To quantify RI events across SF mutant and WT conditions, we utilized IRFinder, a robust tool for detecting and quantifying retained introns from RNA-seq data. Stringent filtering criteria were applied to include only introns with a minimum median junction-spanning read coverage of 10 in each sample, ensuring reliable detection of retention events. Additionally, introns with annotated non-intronic features or flagged with warning tags such as “LowCover” and “MinorIsoform” were excluded to improve specificity. Differential RI (dRI) events were identified by comparing changes in median intron retention levels between SF mutant and WT conditions, using a threshold of |dRI| ≥ 0.58. Statistical significance was assessed via the Wilcoxon rank-sum test, with correction for false discovery rate (FDR < 0.05). This rigorous approach enabled the identification of high-confidence RI events, highlighting patterns of splicing dysregulation in SF mutant conditions.

### Motif density determination and statistical analysis

To assess the role of SRSF1 and hnRNPA0 in splicing regulation, we analyzed the distribution of their putative binding motifs in intronic regions. Using ENCODE RNA-seq datasets from K562 cells, we identified 13,762 introns with loss of RI (LoIR) and 8,788 with gain of RI (GoIR) upon SRSF1 knockdown, based on |log₂ fold change| > 0.58 and FDR < 0.05. Putative motifs for SRSF1 ([AG]GGAG|GGAGGA|[AG]AGGA) and hnRNPA0 (TTATA|TATAG|ATTAGT|ATTTA|AGATAT|AGATATT) were derived from prior literature. To analyze positional motif enrichment, occurrences of the SRSF1 and hnRNPA0 putative motifs were identified using regular expression-based searches, and separately aggregated within the first and last 200 base pairs of intronic regions. Motif densities were further normalized by the total number of introns in each experimental category (GoIR and LoIR). Statistical comparisons of motif densities between GoIR and LoIR introns were conducted using the Mann-Whitney U test.

We applied logistic regression to evaluate the contribution of sequence and structural features to RI changes, incorporating six predictors: SRSF1 motif density, hnRNPA0 motif density, SRSF1/hnRNPA0 ratio, AT content, 3’ splice site strength, and RNA folding free energy. Predictors were scaled using four methods-no scaling, min-max scaling, Z-score scaling, and robust scaling-to assess model stability. The predictive accuracy remained consistent across all scaling methods, with no scaling achieving a slightly higher accuracy (62.14%) compared to min-max scaling (62.13%), Z-score scaling (62.07%), and robust scaling (62.08%). Cross-validation accuracies were also similar, with minimal variation across scaling approaches (e.g., no scaling CV accuracy: 61.25% ± 1.25%, min-max scaling CV accuracy: 61.24% ± 1.31%). VIF analysis confirmed no multicollinearity among the predictors (VIF < 5), ensuring robust feature inclusion.

### Phosphoproteomics sample preparation and phosphopeptide enrichment

Cell pellets were lysed in buffer and subjected to sonication at 4°C using a VialTweeter device (Hielscher Ultrasound Technology; 2 × 1 min) to facilitate protein release. Lysates were cleared by centrifugation at 20,000□×□g for 1 hour at 4°C, and protein concentrations in the resulting supernatant were quantified using the Bio-Rad protein assay (Bio-Rad #5000006). For each sample, 800□µg of protein was diluted in 6□M urea and 100□mM ammonium bicarbonate to a final concentration of 2□µg/µL. Samples were reduced with 10□mM dithiothreitol (DTT) at 56°C for 1 hour and alkylated with 20□mM iodoacetamide (IAA) in the dark at room temperature for 1 hour. Proteins were then processed using a precipitation-based digestion protocol. Briefly, five volumes of pre-chilled precipitation buffer (50% acetone, 50% ethanol, 0.1% acetic acid) were added, and the samples were incubated at -20°C overnight. The precipitated proteins were recovered by centrifugation at 20,000□×□g for 40 minutes at 4°C, washed with ice-cold 100% acetone, and centrifuged again under the same conditions. After removal of residual acetone by SpeedVac, the dried protein pellets were resuspended in 300□µL of 100□mM ammonium bicarbonate and digested overnight at 37°C with sequencing-grade porcine trypsin (Promega) at a 1:20 enzyme-to-substrate ratio. Peptides were then acidified with formic acid and desalted using C18 MacroSpin columns (NEST Group) following the manufacturer’s instructions. Peptide concentrations were measured using a NanoDrop spectrophotometer (Thermo Scientific).

For phosphopeptide enrichment, approximately 500□µg of desalted peptides from each sample was processed using the High-Select™ Fe-NTA Phosphopeptide Enrichment Kit (Thermo Scientific, #A32992) according to the manufacturer’s instructions. Peptides were incubated with resin at room temperature for 30 minutes with gentle shaking, and the mixture was transferred to filter tips (TF-20-L-R-S, Axygen) for low-speed centrifugation (500□×□g, 30 s) to remove unbound peptides. The resin-bound phosphopeptides were washed three times with 200□µL of wash buffer (80% acetonitrile, 0.1% TFA) and twice with 200□µL of water to remove non-specifically bound peptides. Phosphopeptides were eluted twice with 100□µL of elution buffer (50% acetonitrile, 5% ammonium hydroxide), dried by SpeedVac (Thermo Scientific), and reconstituted in MS-compatible buffer. Final phosphopeptide yields were quantified by NanoDrop, and approximately 1.5□µg of enriched material was used for LC-MS/MS analysis.

### DIA-MS Proteomics and Phosphoproteomics Data Analysis

Peptides from total proteome and phosphopeptide-enriched samples were analyzed by data-independent acquisition mass spectrometry (DIA-MS) using an Orbitrap Fusion Lumos Tribrid mass spectrometer (Thermo Scientific) coupled to an EASY-nLC 1200 nanoLC system (Thermo Scientific). Chromatographic separation was performed on a 75□µm × 50□cm C18 column (ReproSil-Pur 120 Å, 1.9□µm particles), with a 120-minute gradient from 6% to 37% buffer B (80% acetonitrile, 0.1% formic acid), followed by a ramp to 100% and re-equilibration. The flow rate was maintained at 300□nL/min with column temperature held at 60°C. DIA acquisition included one MS1 scan (350-1650□m/z, 120,000 resolution) and 40 MS2 scans with variable isolation windows (200-1800□m/z, 30,000 resolution, 28% HCD). MS1 AGC target was set to 2□×□10□ with 100□ms injection time, and MS2 AGC was 5□×□10□ with 50□ms injection time. Spectra were acquired in profile mode with a default charge state of 2. All DIA-MS data were processed using Spectronaut v14 (Biognosys) with the DirectDIA workflow, a spectral-library-free approach. Peptide identification was performed against the Swiss-Prot human protein database (September 2020; 20,375 entries). For the total proteome dataset, variable modifications included methionine oxidation and N-terminal acetylation, and cysteine carbamidomethylation was set as a fixed modification. For the phosphoproteome dataset, phosphorylation on serine, threonine, and tyrosine residues (S/T/Y) was added as an additional variable modification.

Phosphopeptide intensity data were analyzed across wild-type and splicing factor mutant samples to identify differentially phosphorylated sites. For the SRSF2 analysis, phosphopeptide precursor intensities were compared between three SRSF2^WT^ and three SRSF2^P95H^ samples. For the SF3B1 and U2AF1 analyses, comparisons were performed between two wild-type and two mutant samples per condition. All precursor intensities were log₂-transformed, and values below 500 were removed prior to normalization. Mean intensities were calculated for each condition, and fold-change was expressed as log₂(mutant/wild-type). Phosphopeptide precursor ions were collapsed to representative phos.ids by selecting the precursor with the most complete data across runs, or highest summed intensity if tied. Gene symbols, modified sequences, and specific phosphorylation site positions Following up statistical significance was assessed using Student’s t-test, and resulting phosphopeptides were filtered using p-value thresholds (e.g., p < 0.05 or as defined downstream). Only phosphopeptides with confidently localized phosphosites (based on PTM localization scores >0.75 in Spectronaut) were included. were retained to enable downstream mapping to biological pathways and kinase motif enrichment.

### Immunofluorescence

A total of 1.5 × 10□ K562 or CD34+ cells were resuspended in PBS and deposited onto glass slides using a Cytospin (Thermo Scientific) at 1,000 × *g* for 5 minutes. Slides were air-dried for 1 minute and immediately fixed in freshly prepared 4% formaldehyde (Superfrost, Thermofisher) in PBS for 15 minutes at room temperature (RT), followed by three washes with PBS. Cells were permeabilized in ice-cold methanol for 10 minutes and again washed three times in PBS. Blocking was performed using 3% BSA in PBS for 1 hour at RT. Primary antibodies against SRSF1 (rabbit polyclonal, 1:1000) and SC35 (mouse monoclonal, 1:1000) were diluted in blocking buffer and incubated overnight at 4□°C in a humidified chamber. After three washes with PBS, Alexa Fluor-conjugated secondary antibodies (AF555 or AF488, Invitrogen) were applied at a dilution of 1:500 in blocking buffer for 2 hours at RT. Slides were washed five times in PBS and mounted using ProLong™ Gold Antifade Mountant with DAPI (Thermo Fisher Scientific). IF images were acquired using a Leica DM6000 microscope equipped with a DFC390 camera at 63× magnification and LAS AX image acquisition software (Leica). The FIJI (ImageJ) image processing package was used for IF analysis.

### Western blot immunoblotting

Cells were counted and equal cell numbers were harvested and washed in ice-cold PBS. Pellets were resuspended in ice-cold lysis buffer supplemented with Roche protease and phosphatase inhibitor cocktails, flash-frozen (in liquid nitrogen) and thawed on ice three times, then clarified by centrifugation (4°C). Lysates corresponding to equal cell equivalents were mixed with sample buffer, denatured, resolved by SDS-PAGE, and transferred to PVDF. Membranes were blocked in TBST + 1% BSA; both primary (phospho-specific and total) and HRP-conjugated secondary antibodies were diluted in TBST + 1% BSA. Blots were washed in TBST between incubations and developed by chemiluminescence. All steps were performed on ice or at 4 °C unless noted.

### Flow cytometry

Freshly thawed bone-marrow mononuclear cells (BMMNCs) from healthy volunteers and clinically annotated MDS patients (∼10-15 million cells per sample) were washed in PBS + 3% FCS, counted, and incubated with the indicated antibody cocktail in PBS + 3% FCS. After three washes in PBS + 3% FCS, samples were analyzed and flow-sorted according to the gating scheme shown in **Fig. 6A**. For downstream assays, equal numbers of sorted cells were pelleted for each comparison and processed for phosphoprotein immunoblotting exactly as described above (protease + phosphatase inhibitors; flash-freeze/thaw ×3; SDS-PAGE and Western).

### RNA-seq

Total RNA was purified, enzymatically fragmented, and assessed for integrity and size distribution using the Agilent 2100 Bioanalyzer. RNA-seq libraries were generated using the ZapR Mammalian Low Input RNA-seq Kit (Takara Bio) with dual-index primers, following the manufacturer’s instructions. Libraries were sequenced on the Illumina NovaSeq 6000 platform using paired-end 150 bp reads (2 × 150) on an S4 flow cell, achieving an average depth of ∼50 million reads per sample.

### DDR induction in CD34^+^ cells

CD34⁺ hematopoietic progenitors were rapidly thawed at 37 °C, transferred dropwise into pre-warmed SFEM-based medium supplemented with SCF, FLT3L, TPO, GM-CSF, and EPO (manufacturer-recommended concentrations), washed, and plated at 2-5 × 10□ cells/mL to recover overnight (37 °C, 5% CO₂). For UV stress, cells were washed in PBS, placed in a thin layer in uncovered dishes, irradiated with UV-C (254 nm; 5 (low dose) or 10 (high dose) mJ/cm²), returned to complete medium, and harvested 4 h post-irradiation. For chemical DDR induction, cells were treated in complete medium with camptothecin (1 µM), etoposide (10 µM), or hydroxyurea (2 mM) (final DMSO ≤ 0.1%) and harvested 36 h later. After treatment, equal numbers of cells per condition were collected for phosphoprotein immunoblotting (as described above) or fixed/permeabilized for imaging. For RNA-seq profiling, DDR-induced and matched physiological control cells received a 4 h pulse of 4-thiouridine (4sU) immediately prior to harvest; 4sU-labeled RNA was then selectively biotinylated and enriched on streptavidin beads before library preparation.

### Methylcellulose colony forming unit assay

BMMNCs were isolated using Ficoll-Hypaque density gradient centrifugation (Histopaque-1077, Sigma Aldrich) and plated in methylcellulose medium (Methocult GF#H4435, Stem Cell Technologies) following the manufacturer’s instructions. For rescue experiments, the BMMNCs were transduced with lentiviral vectors expressing human or *E. coli* RNaseH1 by spinfection (1800 rpm, 90 min, 37°C) in the presence of 8 µg/mL polybrene. Following a 6-hour recovery in IMDM supplemented with 10% FBS at 37°C, stably transduced cells were replated in methylcellulose medium as above and incubated for 14 days. After 14 days of incubation, erythroid and myeloid colonies were manually counted using a STEMgrid-6 (StemCell Technologies, #27000), and representative colony images were captured with a Keyence BZX-800 inverted fluorescence microscope. After colony imaging, cells were harvested for downstream analyses, including western blotting (for phosphoprotein changes), low-input RNA-seq (to assess alternative splicing), and Sanger sequencing of cDNA (to confirm mutant clone representation).

## Data Accession

Patient RNA-seq data from Hershberger et al study^13^ (splicing-factor-mutant myeloid disorders, n=395; controls, n=64) were analyzed using author-provided processed outputs. These processed files were supplied under a collaboration agreement and may be shared by the corresponding authors upon reasonable request and with permission from the data owners. The underlying raw sequencing files are held by the data owners and are not publicly available due to participant consent and data-protection regulations. Access requests should be directed to Dr. Torsten Haferlach and will require a data-use agreement.

## Author Contributions

PCB and MMP conceived and designed the study. PCB and RR performed the microscopy, Western blotting, and cell culture assays. CA, WL, and YL prepared samples for mass spectrometry, processed the data, and carried out the initial analyses; PCB generated the corresponding figures. FF, GT, and MDP provided MDS patient samples for flow cytometry and Western blot analyses. SH, FB, and TH provided processed RNA-seq outputs from patient and healthy volunteer cohorts for downstream analyses. PCB and MMP wrote and edited the manuscript. MMP supervised the overall study.

## Acknowledgements

This work was funded in part by the National Institutes of Health (HL167071 and CA270656 to MMP; K08 1K08DK140636 to PCB), a pilot award from Yale Center for Gastrointestinal Cancers (CGIC) to PCB. We thank the Yale Center for Genome Analysis (YCGA), Yale Center for Research Computing (YCRC), and Yale Core Center of Excellence in Hematology (YCCEH, supported by NIDDK U54 DK106857) for the core resources used. Some of the patient samples used in this study were provided by The Yale Hematology Tissue Bank. We thank Can Akpinaroglu for contributions to mass spectrometry sample processing.

## Diversity Statement

We support inclusive, diverse, and equitable conduct of research.

## Figure legends

**Supplemental Figure 1:** (A) Splicing factor mutation landscape in 7,554 MDS patients (MSKCC 2020 and IWG 2022) from Cbioportal.^106^ The oncoprint (top) displays mutation statuses for *SF3B1*, *U2AF1*, *SRSF2*, and *ZRSR2*. Each horizontal bar represents an individual gene across the patient cohort. Green bars indicate mutations (darker green: putative driver mutations; lighter green: mutations of unknown significance), and gray indicates no alterations. The table (bottom) shows pairwise analyses of co-occurrence versus mutual exclusivity. Negative Log2 Odds Ratios and corresponding q-values < 0.05 suggest significant mutual exclusivity between the indicated genes. (B) Box-and-whisker plot of log2-transformed median TPM values for each condition, with colors corresponding to the presence or absence of an epigenetic modifier (EM). Boxes show the interquartile range (IQR), whiskers extend to 1.5*IQR, and circles represent outliers. (C) The diagram shows the total RI events (10,645) with a subset (4,151) identified as IRFinder-based RI events having good coverage and no overlapping features. Events excluded due to lack of annotation, low coverage, or overlapping non-intronic features account for 6,494. (D) Stacked bar plot showing the distribution of alternative splicing events across seven conditions and five AS types, with bars divided into commonly significant events (blue), unique significant events (not significant across all seven categories; orange), and non-significant events (gray). Percentages indicate the proportion of events within each condition and AS type. (E) Bar plot comparing observed and expected counts across five AS event types (MXE, SE, RI, A3, A5). Percent differences between observed and expected values are shown above the observed bars. The observed values represent the actual counts of significant AS events for each AS type. The expected values are derived from the chi-squared test, assuming a null hypothesis where the proportion of significant events matches the overall distribution of AS events. The chi-squared statistic and p-value evaluate the deviation between observed and expected distributions. RI events show the highest positive deviation, emphasizing their significant contribution. (F) Heatmap of RI splicing event differences clustered by condition. Rows represent events, and columns represent samples for splicing factor mutations with or without EM, as well as WT. (G) Heatmap showing hierarchical clustering of RI splicing event changes across conditions from data published by Pellagatti et al. Columns represent samples grouped by SF mutations and controls, with annotations indicating specific conditions. (H) t-SNE analysis of RI profiles in SF mutants and hematopoietic lineages. As in Fig. 1E, SF3B1-, SRSF2-, and U2AF1-mutant samples form distinct clusters. Convex hulls and centroids are shown for each group to illustrate cluster boundaries and central positions.

**Supplemental Figure 2:** (A-D) Comparisons of Intron Characteristics Across Categories: Gain of Intron Retention (GoIR), Loss of Intron Retention (LoIR), and No Change in Intron Retention (ncIR).(A) Log10 Transformed Intron Lengths: Boxplots show the distribution of log10-transformed lengths for introns in each category.(B) Intronic GC Content: Comparison of GC content (percentage) across intron categories reveals higher GC content in GoIR introns compared to LoIR and ncIR introns.(C) Free Energy: The distribution of free energy (kcal/mol) shows that GoIR introns exhibit significantly lower free energy, indicative of potentially stable secondary structures, compared to LoIR and ncIR introns. (D) Conservation Scores: Boxplots show the conservation scores for introns across the three categories. (E) Plots as in Fig 2G. Rows: U2AF1_mut_ (top) and SRSF2_mut_ (bottom). Columns: All introns, Short+mid (lower two length tertiles combined), and Long (upper tertile). Colors: LoIR (blue) and GoIR (salmon). Shaded histograms with overlaid KDEs are shown; the dashed line marks 0. Text in each panel reports the one-sided Wilcoxon p-value testing median < 0 for each group. Overall, the left-shift (negative Δ) is most evident in short+mid introns, with minimal shift in long introns. (F) Multivariable logistic regression (HC3 robust SEs) of Pr(Δ (residual, sign-aligned) < 0). Predictors z-scored; values shown are OR per 1-SD increase with 95% CI, Wald p, and Benjamini-Hochberg q. N=1115 introns; AUC=0.599. An interaction model was also fit; no interaction term was FDR-significant (AUC=0.603). (G) Panels show the distribution of Δlog-slope (SF_mutant_ − WT) stratified by intron length. Δlog-slope values are residualized vs WT and sign-aligned to WT**, so** negative values indicate a flatter-than-expected 3′→5′ coverage gradient in the mutant. LoIR (blue) and GoIR (salmon). Shaded histograms with overlaid KDEs are shown; the dashed line marks 0. Text in each panel reports the one-sided Wilcoxon p-value testing median < 0 for each group. Overall, the left-shift (negative Δ) is most evident in short+mid introns, with minimal shift in long introns.

**Supplemental Figure 3:** (A) Hierarchical clustering dendrogram of 3,518 significant RI events, identified across three splicing factor mutation categories, as detailed in (Fig. 2A-B). Each branch represents an individual shRNA knockdown condition from ENCODE datasets, encompassing both HepG2 and K562 cell lines. Branches corresponding to SRSF1 knockdown and mutation-induced RI alterations highlighted in blue and green respectively. (B) List of RNA binding protein shRNA knockdown conditions obtained from K562 ENCODE data. Heatmap illustrating the hierarchical clustering of delta RI (dRI) changes involving (C) skipped exon (SE) and (D) other (MXE, A5, A3) events across multiple shRNA knockdown conditions in K562 cells. Each row represents an individual RI event, while each column corresponds to a specific knockdown condition. The color intensity reflects the magnitude of dRI changes, with red indicating increased retention and blue indicating decreased retention. (E) Histograms depicting the distribution of various putative SRSF1 and hnRNPA0 motif densities within 200 bp intronic regions flanking the 5’ and 3’ splice sites (5’SS -> 200bp followed by 200bp -> 3’SS) in LoIR_SRSF1KD_ and GoIR_SRSF1KD_ intron categories. Comparison of (F) 3’ splice site score strength, (G) GC content of 200bp intronic region flanking the 5’ and 3’SS, and (H) RNA free energy at 3’SS, between GoIR_SRSF1KD_ and LoIR_SRSF1KD_ category introns. (I) Logistic regression analysis of intron retention predictors. The table summarizes regression coefficients, standard errors, z-scores, p-values, and 95% confidence intervals for six predictors: SRSF1 motif density, hnRNPA0 motif density, SRSF1/hnRNPA0 motif ratio, AT content, 3’ splice site strength, and RNA folding free energy.

**Supplemental Figure 4:** (A) Volcano plots illustrating differential gene expression in SF mutant versus WT conditions, based on data from Pellagatti et al. Each point represents a gene, with its x-axis value indicating the Log2FC and its y-axis value indicating the -log10 of FDR. Highlighted genes, including SRPK1, CLK1, and SRSF1, are annotated in the plots to indicate their positions relative to the thresholds.(B) Correlation heatmap of Pearson correlation coefficients for gene expression profiles across biological replicates of SRSF2^P95H^ and SRSF2^WT^ conditions. Higher correlation values (closer to 1.0, shown in red) indicate greater similarity in gene expression patterns within and across conditions. (C) Principal Component Analysis (PCA) of replicates for SRSF2^P95H^ (blue) and SRSF2^WT^ (orange) conditions. PCA reveals clear separation between mutant and wild-type replicates along PC1 (40.59% of variance) and PC2 (23.09% of variance), indicating distinct transcriptional profiles. (D) Heatmap showing hierarchical clustering of differentially expressed P-sites (Z-scores) between SRSF2^P95H^ mutant vs SRSF2^WT^ samples. Red indicates upregulation, and blue indicates downregulation relative to the mean expression. Associated biological processes significantly enriched in each cluster (log10P > 10) are annotated. (E) Schematic of SRSF1 domains and phosphorylation analysis. The RS1 domain (amino acids 197-226) contains phosphorylation sites S199 and S201. Box-whisker plot shows log2 intensity of total SRSF1 in WT (purple) vs. mutant (orange) conditions for SF3B1 and U2AF1.

**Supplemental Figure 5:** (A) Volcano plot of AKT-target phosphosites in SF3B1^K700E^ vs SF3B1^WT^ (left panel) and U2AF1^S34F^ vs U2AF1^WT^ (right panel). Differentially phosphorylated sites are highlighted with significant fold change (L2FC±0.58 and FDR < 0.1.) Key AKT-regulated upregulated (red) and downregulated (blue) phosphosites are annotated. (B) Box plot showing the relative log2-transformed intensity of mTOR Ser-2481 phosphorylation in SRSF2^WT^ and SRSF2^P95H^ mutant conditions. The data highlights a significant reduction in mTOR phosphorylation in the mutant compared to the wild type (FDR = 0.018). Error bars represent the range of replicates. (C) Western blot analysis and quantification of signaling and splicing proteins in SF3B1^WT^, SF3B1^K700E^, U2AF1^WT^, and U2AF1^S34F^ cells. Proteins analyzed include RAPTOR (total and phosphorylated), SRPK1, AKT1 (total and phosphorylated), and SRSF1 (total and phosphorylated). Quantification of image intensity (a.u.) for protein ratios such as P-SRPK1/SRPK1, P-AKT1/AKT1, and P-RAPTOR/RAPTOR reveals significant changes between wild-type (violet bars) and mutant (orange bars) cells, with p-values indicating statistical significance. D) Heat-map of ΔPSI values for RI events that shift in the same direction in SF-mutant cells and at least one DNA-damage treatment (UV low, UV high, camptothecin, etoposide, hydroxyurea). Green indicates decreased retention; red indicates increased retention. (E) Overlap size of RI events between SF mutant state and DDR stress treatments. Heatmap shows the number of introns altered in both SF mutant state and each treatment across increasing magnitude thresholds (|Δ|). (F) Directional concordance of RI events with SF mutant state. Heatmap shows the percentage (and number) of overlapping introns that change in the same direction in SF mutant state and each treatment, across magnitude thresholds. (G) Venn diagram showing the number of significant RI events with non-zero ΔPSI in SF mutant cells that also exhibit concordant directionality (i.e., same sign of ΔPSI) in response to UV-high dose or camptothecin treatment. Events were considered concordant if the sign of RI change matched that observed in the SF mutant. The diagram illustrates RI events unique to UV concordance (1270), camptothecin concordance (1201), concordant with both treatments (1106), and SF mutant-specific events without concordance to either treatment (1341). (H) R-Loop Levels in Splicing Factor Mutant and Wild-Type Cells. R-loop levels were measured using the S9.6 antibody in wild-type and mutant cells under varying conditions, including RNH1 overexpression. Dot blot results reveal increased R-loop accumulation in SF mutant cells compared to wild-type cells, with a notable reduction upon RNH1 overexpression. Methylene blue staining serves as a loading control, confirming equal input amounts for the assays. (I) Rescue of Splicing and Signaling Defects by RNAseH1 Overexpression. Western blot analysis of splicing and signaling proteins in SF3B1^WT^, SF3B1^K700E^, U2AF1^WT^, and U2AF1^S34F^ cells, with either empty vector (EV) or RNH1 overexpression. Quantification of protein ratios, including P-SRPK1/SRPK1, P-AKT1/AKT1, and P-RAPTOR/RAPTOR, demonstrates that RNH1 overexpression partially rescues splicing and signaling defects observed in SF mutant cells. Image intensity (a.u.) quantifications further highlight these trends.

**Supplemental Figure 6:** (A) Top enriched pathways from pathway analysis in the three SF mutant (vs WT) conditions. Bar plots depict −log10(p-value) for each pathway, categorized by z-score (positive, negative, or neutral) to indicate predicted activation or inhibition. Pathways without available activity patterns are shown in gray. The orange line represents the significance threshold. (B) Bar plots showing the proportion of genes associated with upregulated, downregulated, and unchanged activity across key signaling pathways in SF mutant versus WT conditions. Pathways are categorized based on their relevance to observed transcriptomic changes, with annotations indicating the percentage distribution of gene activity and the number of associated genes. Pathways with no overlap in the dataset are shown in gray. (C) Representative example showing chromatograms of PCR amplicons around the SF3B1 K666 locus. Patient-derived SF3B1K666Q MDS CD34+ cells were isolated and grown in methylcellulose medium for 14 days after being transduced with lentiviral constructs overexpressing human RNH1, E.coli RNH1, and empty vector. DNA was isolated from the cell colonies at 14 days for the respective conditions and PCR performed on the extracted DNA to amplify the region around the SF3B1 K666 locus. Highlighted in blue is the K666 locus, showing the presence of CAG mutation (K→Q). (D) Hierarchical clustering of alternative-splicing ΔPSI profiles across four event classes (RI, SE, A3, A5). For each class, we computed ΔPSI (OE - EV) for human and E. coli RNase H1 overexpression (blue), and included published knockdowns of hnRNPA0/hnRNPAB (green) and SRSF1 (red). Clustering was performed by Ward linkage on Euclidean distances. In all event types, both RNH1-OE signatures co-cluster with hnRNP knockdowns, whereas SRSF1_KD segregates onto a separate branch.

