## Supplementary figures for "A Unifying Mechanism for Shared Splicing Aberrations in Splicing Factor Mutant Cancers"

### SUPPLEMENTAL FIGURE 1

A

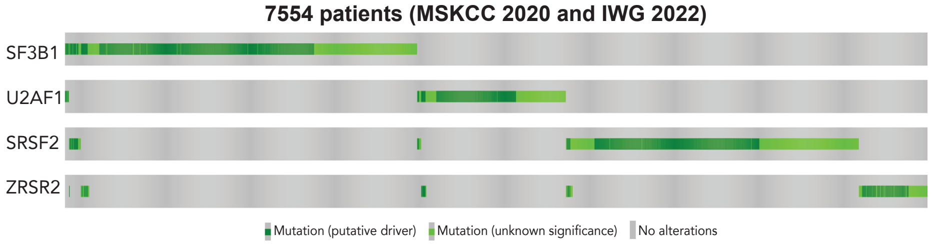


| Mutation A | Mutation B | Log2 Odds Ratio | q-Value | Tendency |
|---|---|---|---|---|
| **SF3B1** | SRSF2 | -2.330 | <0.001 | Mutual exclusivity |
| **SF3B1** | U2AF1 | -2.791 | <0.001 | Mutual exclusivity |
| **U2AF1** | SRSF2 | -2.566 | <0.001 | Mutual exclusivity |
| **SRSF2** | ZRSR2 | -0.782 | 0.014 | Mutual exclusivity |
| **SF3B1** | ZRSR2 | -0.700 | 0.014 | Mutual exclusivity |

B

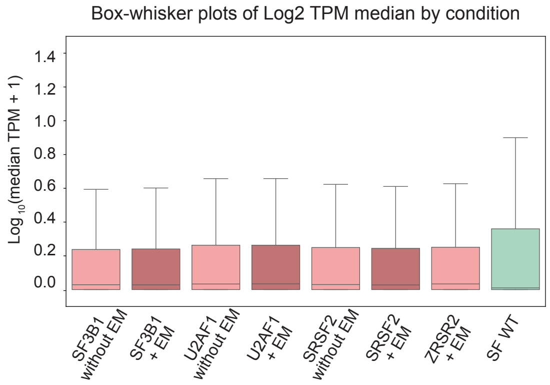


C

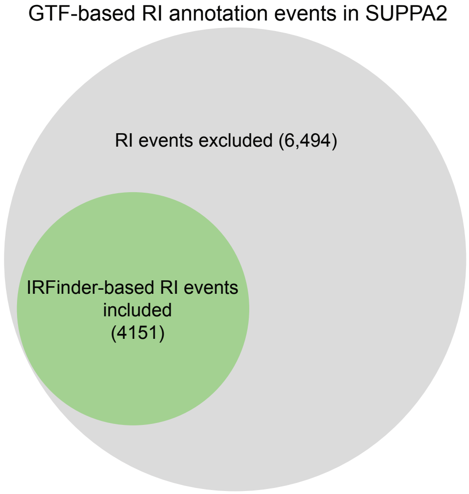


D

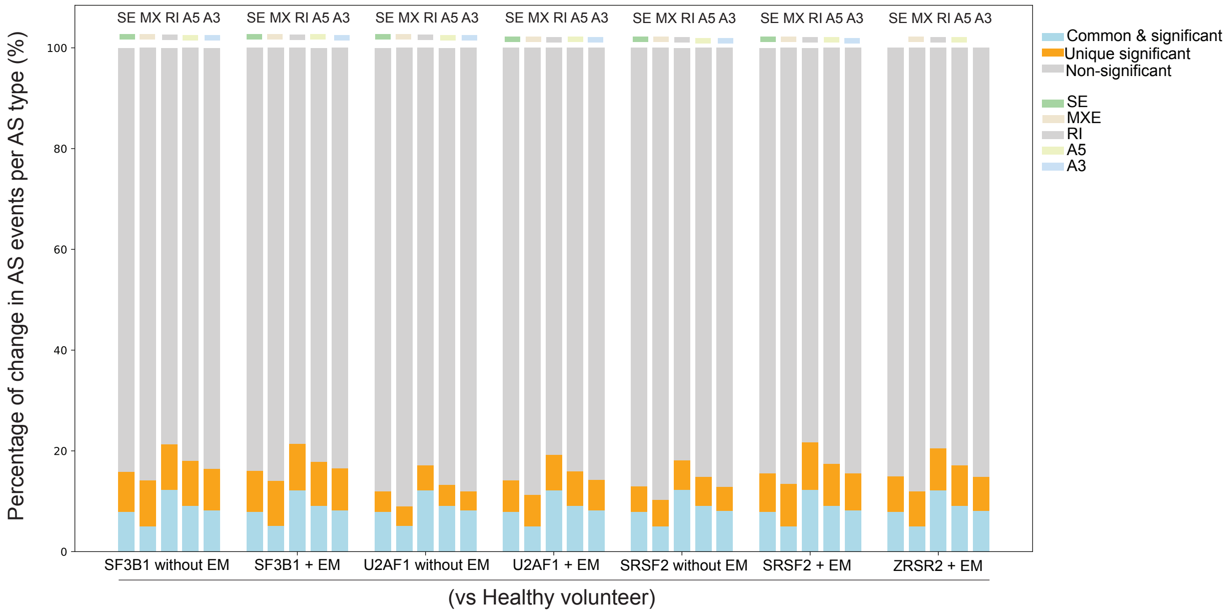


E

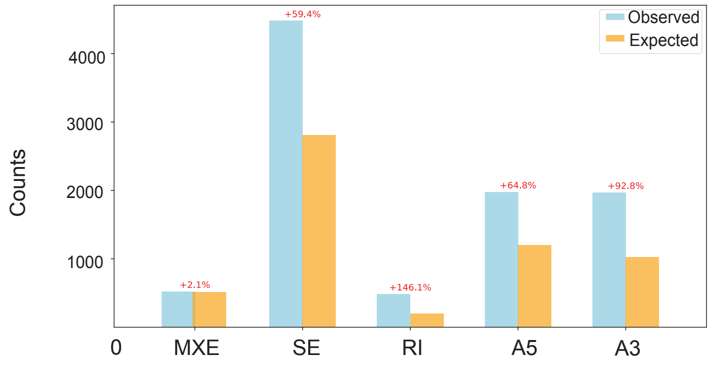

### SUPPLEMENTAL FIGURE 1

F

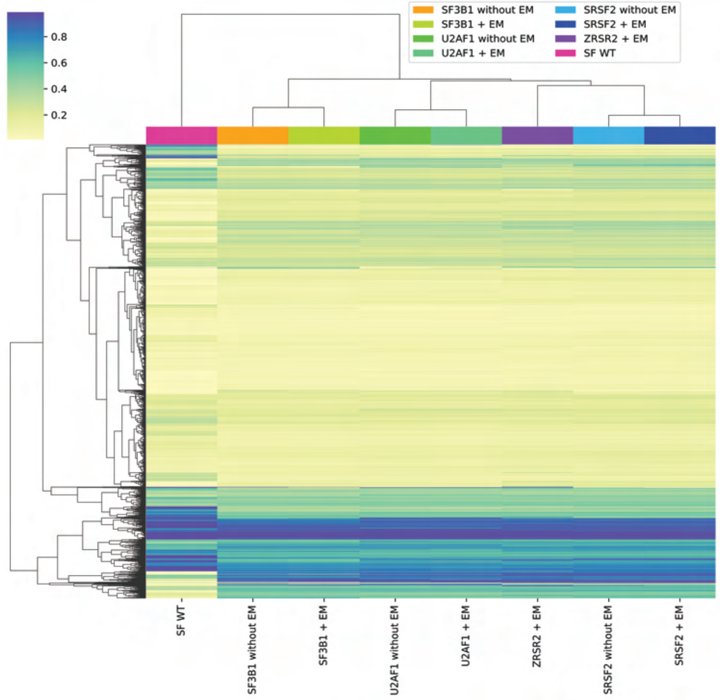


G

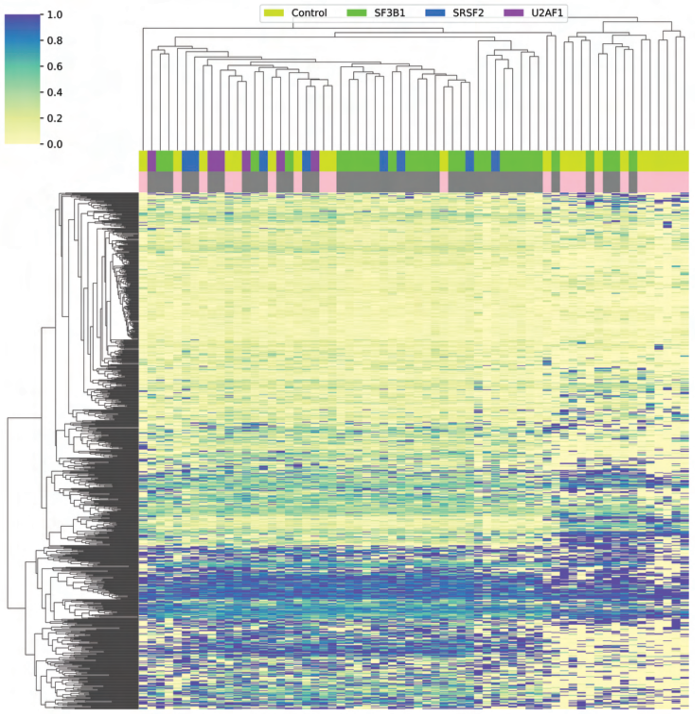


H

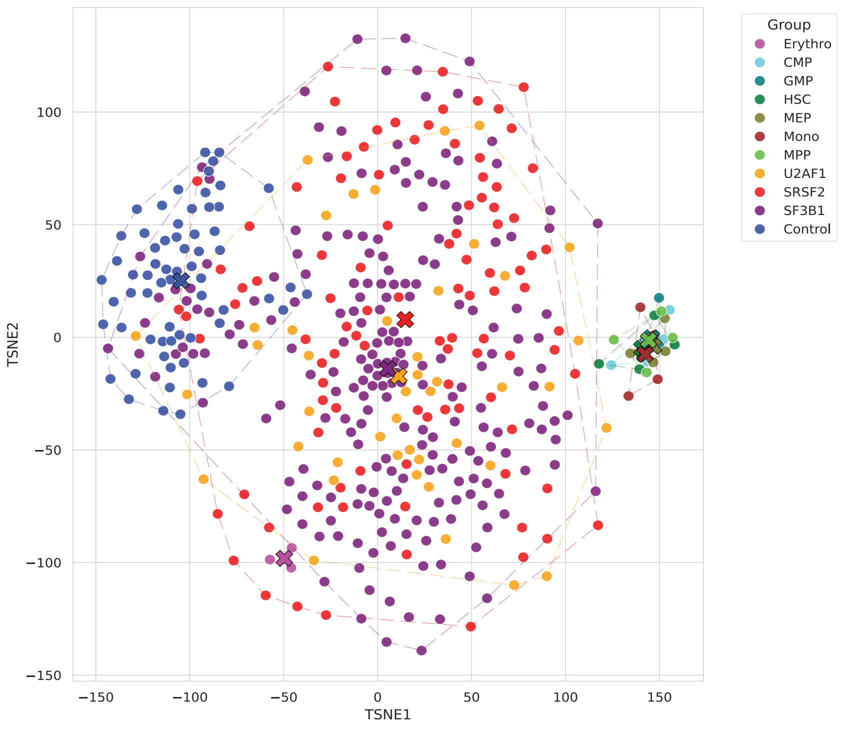

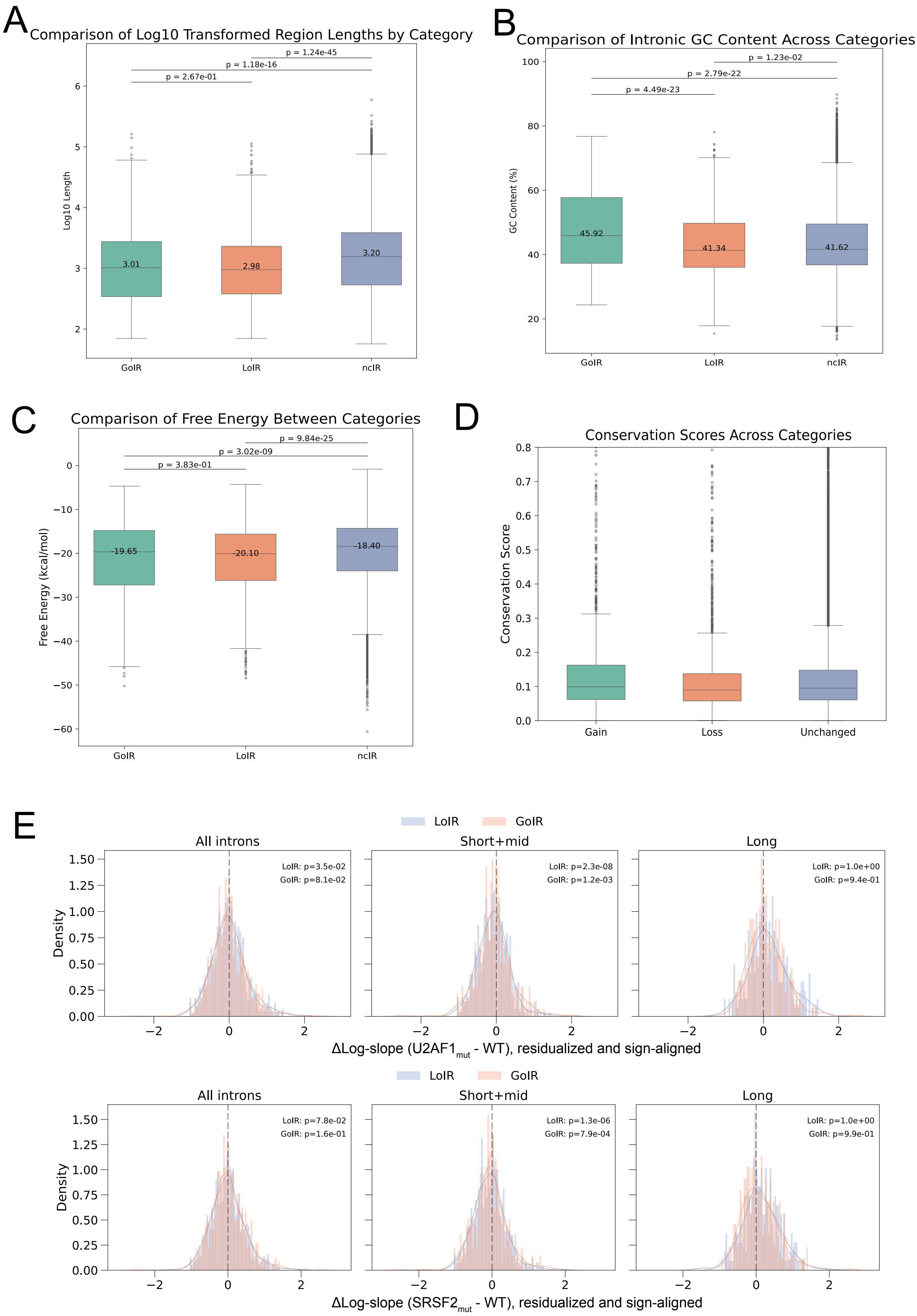
A
Comparison of Log10 Transformed Region Lengths by Category
p = 1.24e-45
p = 1.18e-16
p = 2.67e-01
Log10 Length
3.01
2.98
3.20
GoIR
LoIR
ncIR
B
Comparison of Intronic GC Content Across Categories
p = 1.23e-02
p = 2.79e-22
p = 4.49e-23
GC Content (%)
45.92
41.34
41.62
GoIR
LoIR
ncIR
C
Comparison of Free Energy Between Categories
p = 9.84e-25
p = 3.02e-09
p = 3.83e-01
Free Energy (kcal/mol)
-19.65
-20.10
-18.40
GoIR
LoIR
ncIR
D
Conservation Scores Across Categories
Conservation Score
Gain
Loss
Unchanged
E
LoIR
GoIR
All introns
Short+mid
Long
Density
LoIR: p=3.5e-02
GoIR: p=8.1e-02
LoIR: p=2.3e-08
GoIR: p=1.2e-03
LoIR: p=1.0e+00
GoIR: p=9.4e-01
ΔLog-slope (U2AF1mut - WT), residualized and sign-aligned
LoIR
GoIR
All introns
Short+mid
Long
Density
LoIR: p=7.8e-02
GoIR: p=1.6e-01
LoIR: p=1.3e-06
GoIR: p=7.9e-04
LoIR: p=1.0e+00
GoIR: p=9.9e-01
ΔLog-slope (SRSF2mut - WT), residualized and sign-aligned

F

=== Multivariable logistic (MAIN, HC3 robust) ===

N=1115   AUC=0.599

| Predictor | OR | CI_lo | CI_hi | p | q_BH |
|---|---|---|---|---|---|
| Length (z) | 0.414284 | 0.296078 | 0.579684 | 2.726622e-07 | 0.000002 |
| FE (kcal/mol, z) | 1.116704 | 0.962075 | 1.296186 | 1.466239e-01 | 0.439872 |
| IR group (LoIR vs GoIR) | 0.898766 | 0.685542 | 1.178308 | 4.398312e-01 | 0.859928 |
| 5′SS score (z) | 0.978747 | 0.869117 | 1.102205 | 7.230109e-01 | 0.859928 |
| GC% (z) | 1.022191 | 0.879276 | 1.188334 | 7.751578e-01 | 0.859928 |
| 3′SS score (z) | 0.988848 | 0.873040 | 1.120018 | 8.599280e-01 | 0.859928 |

G

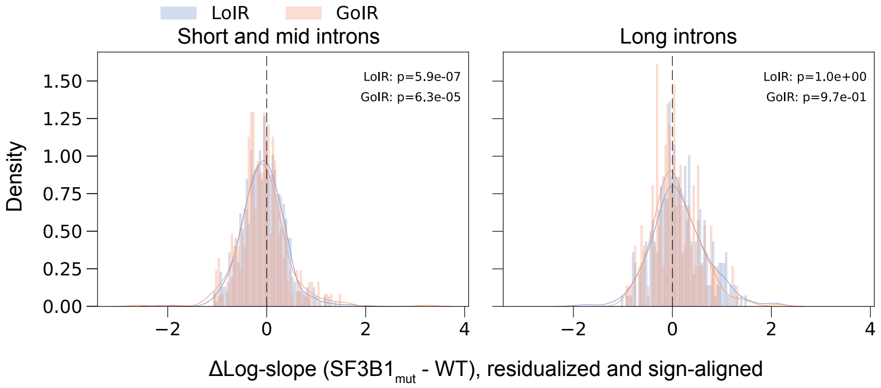

### SUPPLEMENTAL FIGURE 3

A

| Splicing Regulators | RNA Stability and Decay Factors | Nuclear RNA Processing and Transport | Translation Regulators | Chromatin and Transcription Regulation | Cytoplasmic RNA Regulators | RNA Helicases | Ribosome-Associated Proteins | Stress Granule and Processing Body Components | RNA-Binding Miscellaneous |
|---|---|---|---|---|---|---|---|---|---|
| HNRNPA0 | UPF1 | ALYREF | EIF2S2 | AHCTF1 | G3BP1 | DDX5 | RPL5 | AQR | METTL3 |
| HNRNPA1 | UPF2 | CPSF1 | EIF3A | BCCIP | G3BP2 | DDX6 | RPL11 | BUD13 | METTL14 |
| HNRNPA2B1 | UPF3 | CPSF2 | EIF3B | BCLAF1 | FMR1 | DDX17 | RPL18 | LIN28B | MOV10 |
| HNRNPD | EXOSC1 | CPSF3 | EIF4A1 | DNAJC21 | FUS | DDX18 | RPSA | PCGF2 | CDC5L |
| HNRNPF | EXOSC2 | CPSF4 | EIF4G2 | HNRNPK | ELAVL1 | DDX21 | RPS27 | PCGF6 | RNASEH2B |
| HNRNPH1 | EXOSC3 | CPSF6 | EIF4H | HNRNPL | ILF2 | DDX23 | RPL35A | ELF2 | CPSF2 |
| HNRNPK | EXOSC5 | PRPF3 | LARP7 | KHSRP | ILF3 | DDX39A | FAU | | RBM8A |
| HNRNPL | EXOSC7 | PRPF4 | LARP4 | MATR3 | YBX1 | DDX39B | | | |
| HNRNPM | EXOSC8 | PRPF6 | MAGOH | TAF15 | SBP | DDX41 | | | |
| HNRNPU | EXOSC9 | PRPF8 | RPS3 | NONO | HUR | DDX47 | | | |
| HNRNPUL1 | NCBP1 | QKI | RPS3A | RBM10 | | DDX52 | | | |
| SRSF1 | NCBP2 | SNRNP70 | RPS7 | TROVE2 | | DDX55 | | | |
| SRSF2 | TBRG4 | SMN1 | RPS8 | | | | | | |
| SRSF3 | | NPM1 | RPS9 | | | | | | |
| SRSF4 | | PABPC1 | TIA1 | | | | | | |
| SRSF5 | | PABPC3 | TIAL1 | | | | | | |
| SRSF6 | | | | | | | | | |
| SRSF7 | | | | | | | | | |
| U2AF1 | | | | | | | | | |
| U2AF2 | | | | | | | | | |
| TARDBP | | | | | | | | | |
| RBM5 | | | | | | | | | |
| RBM6 | | | | | | | | | |
| RBM39 | | | | | | | | | |
| RBM15 | | | | | | | | | |

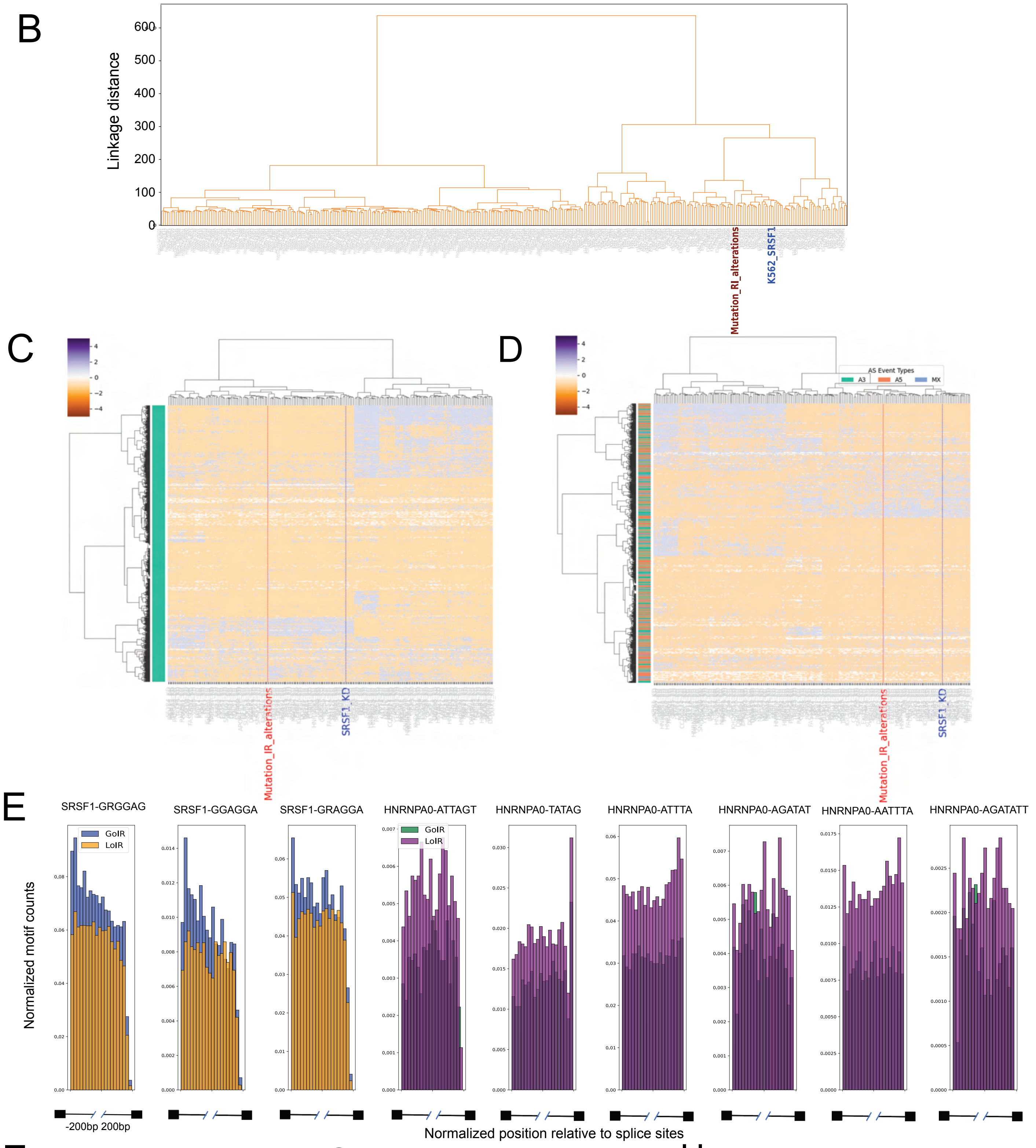


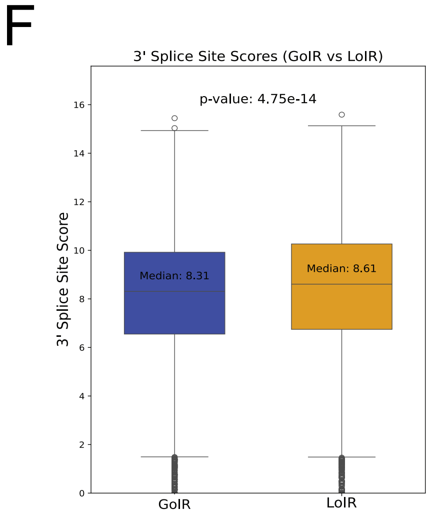


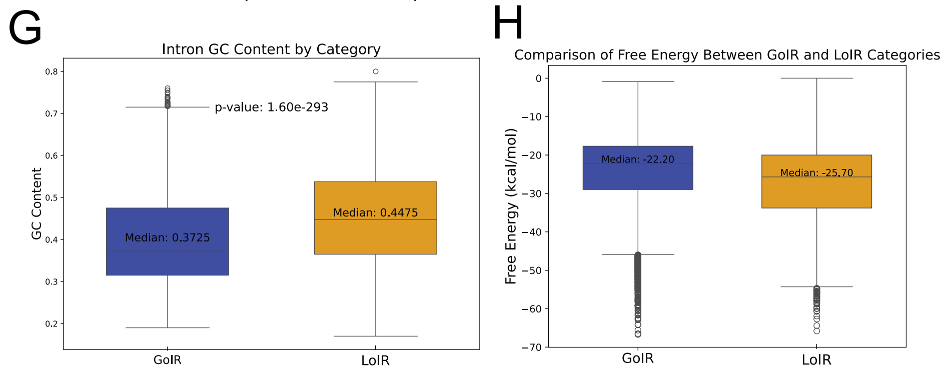


I

| | coef | std err | z | P>\|z\| | [0.025 | 0.975] |
|---|---|---|---|---|---|---|
| const | 0.6007 | 0.264 | 2.271 | 0.023 | 0.082 | 1.119 |
| SRSF1_MOTIF | 0.8118 | 0.174 | 4.657 | 0.000 | 0.470 | 1.153 |
| HNRNPAO_MOTIF | −1.4158 | 0.157 | −9.019 | 0.000 | −1.723 | −1.108 |
| SRSF1/HNRNPA0 | −1.3043 | 0.208 | −6.266 | 0.000 | −1.712 | −0.896 |
| AT_CONTENT | −2.3071 | 0.141 | −16.316 | 0.000 | −2.584 | −2.030 |
| 3SS_SCORE | 0.5044 | 0.282 | 1.786 | 0.074 | −0.049 | 1.058 |
| FREE_ENERGY | 0.2200 | 0.135 | 1.630 | 0.103 | −0.045 | 0.485 |

### SUPPLEMENTAL FIGURE 4

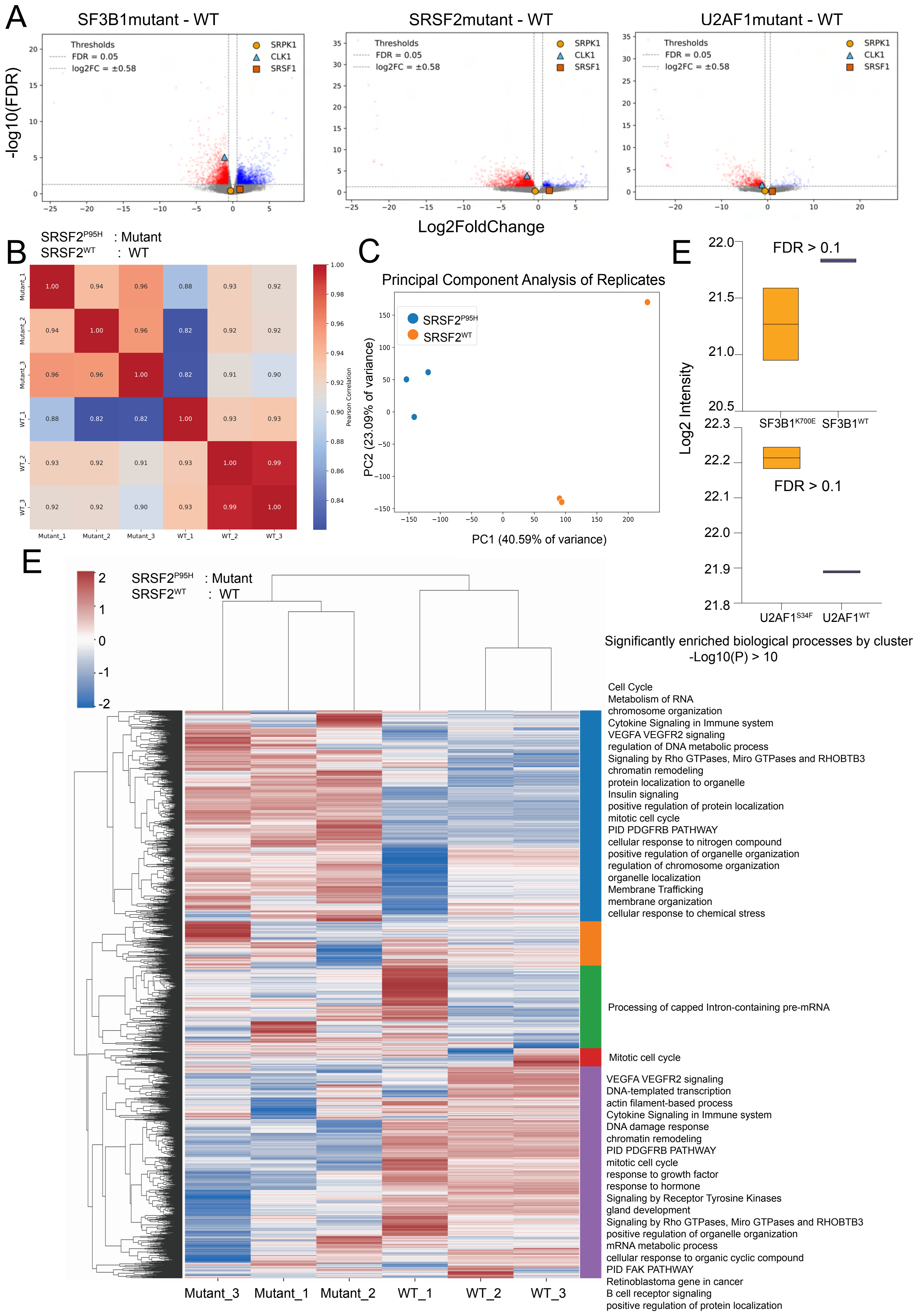

### SUPPLEMENTAL FIGURE 5

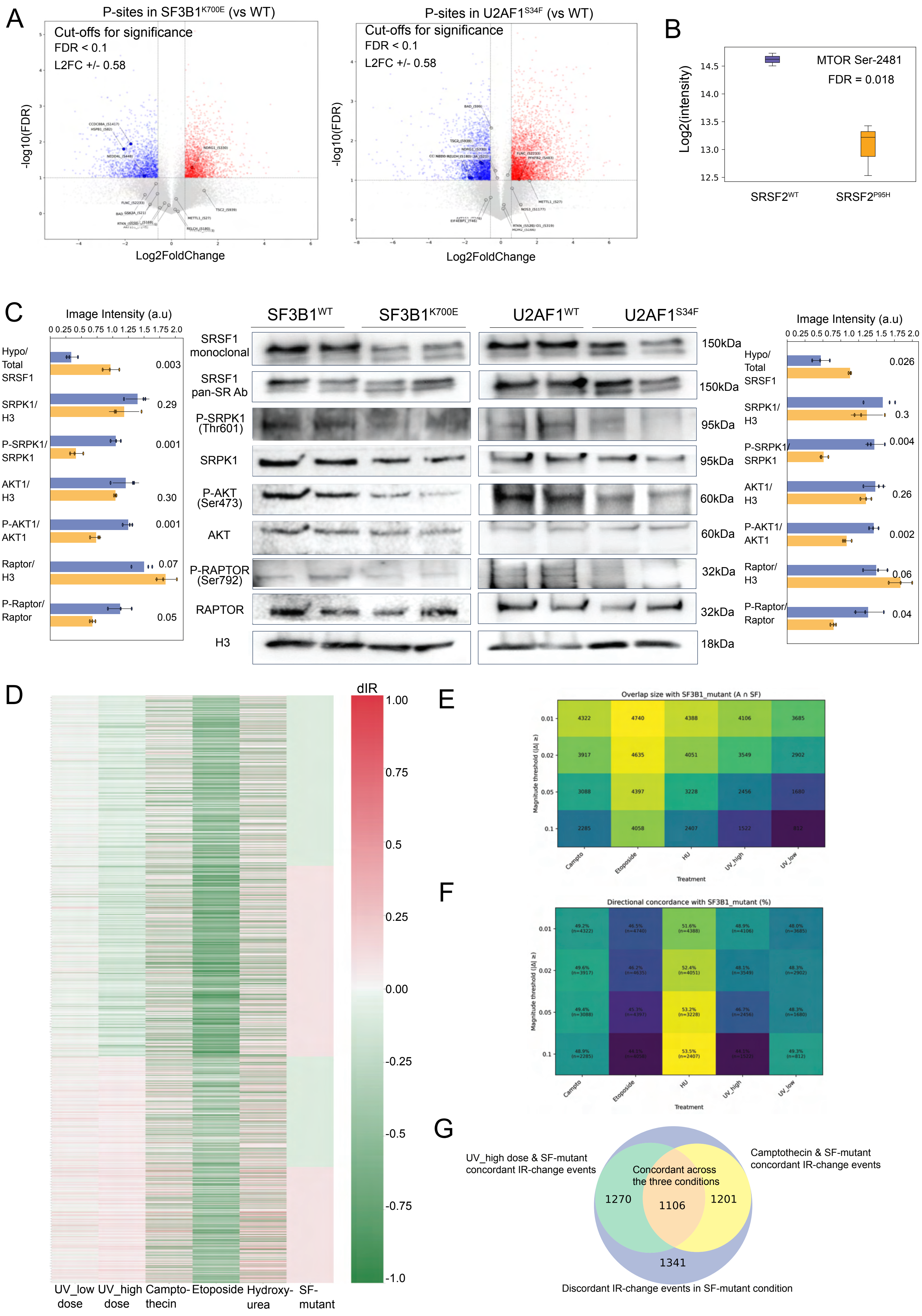

### SUPPLEMENTAL FIGURE 5

F

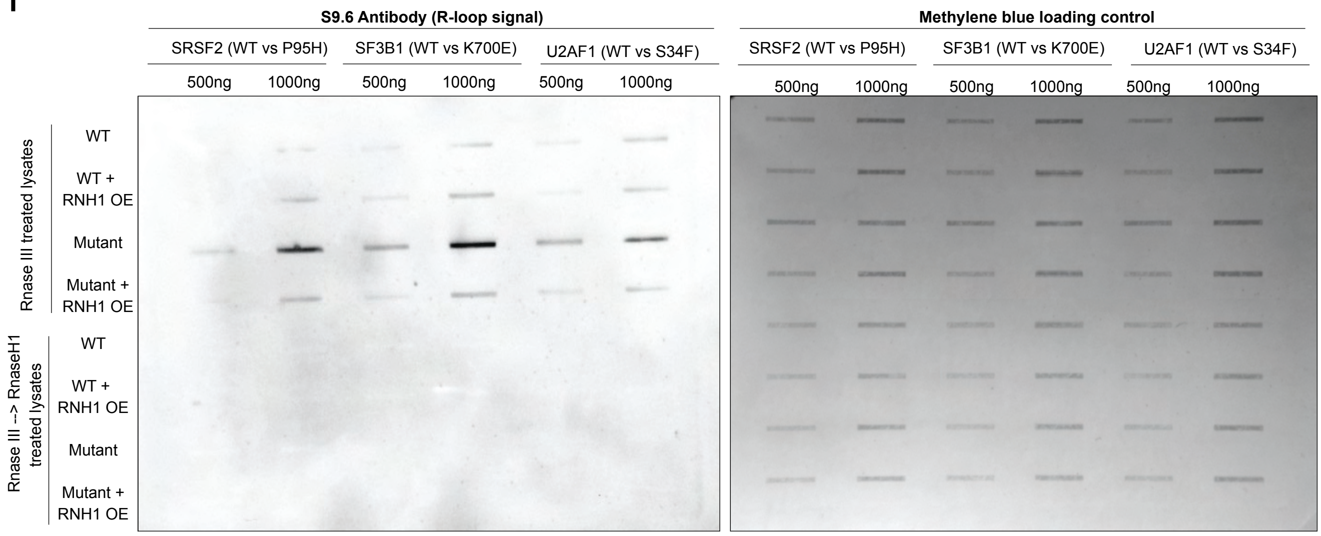


G

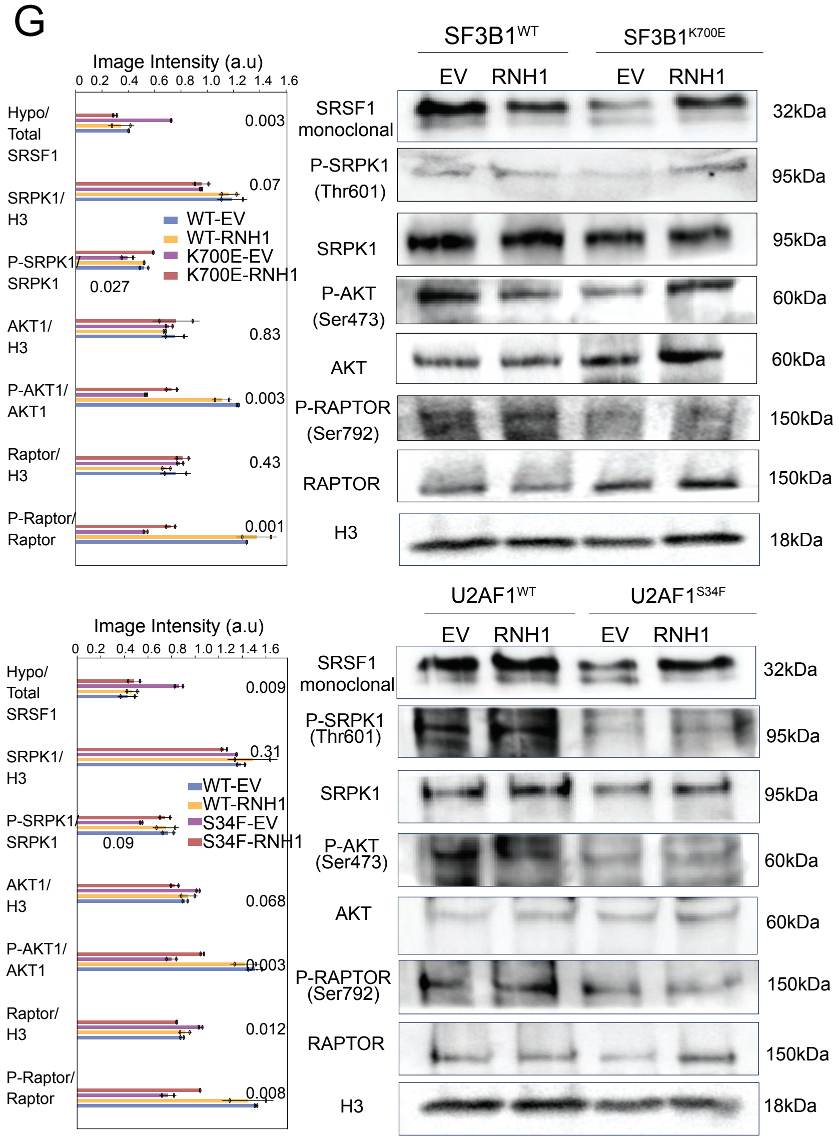

### SUPPLEMENTAL FIGURE 6

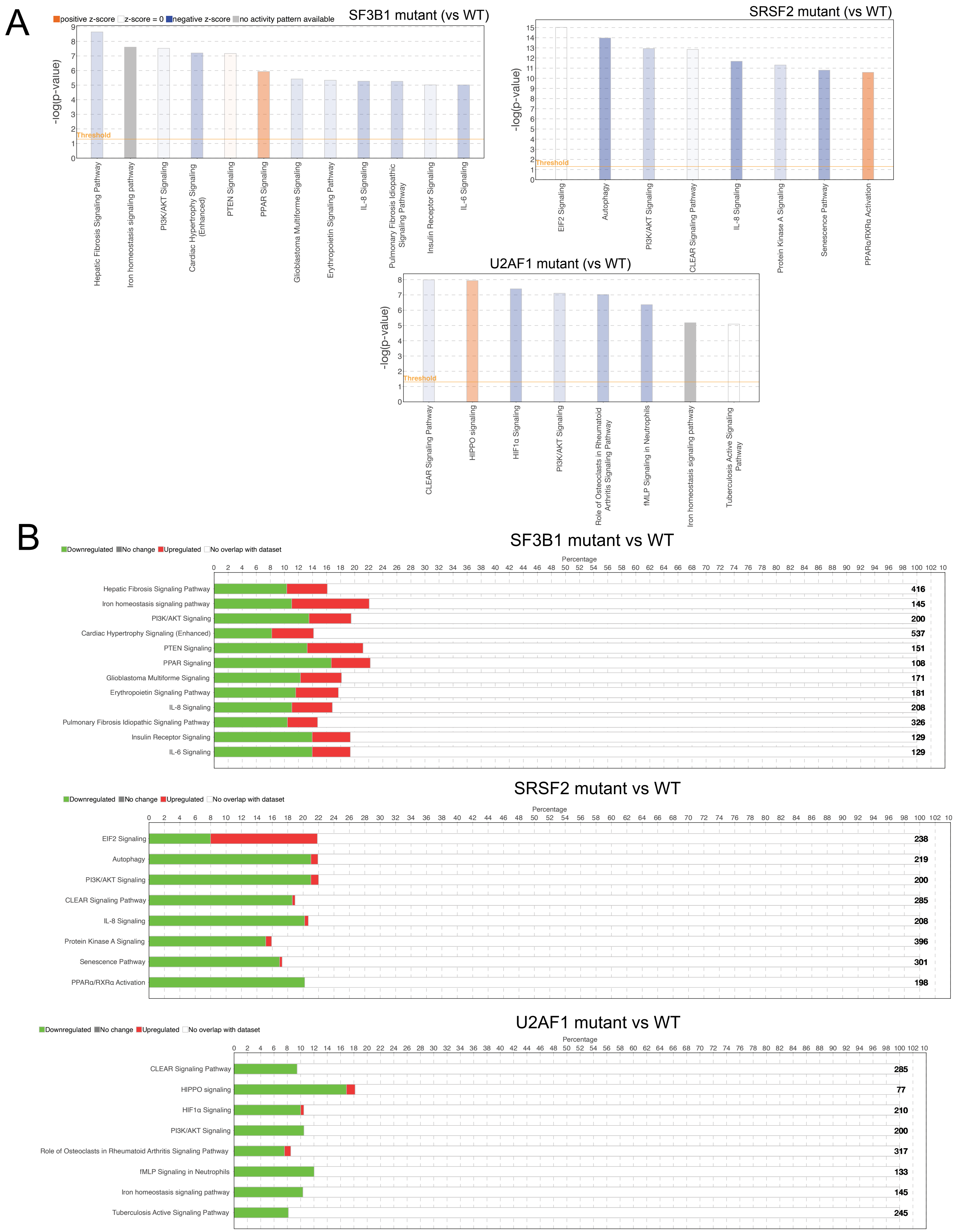

C

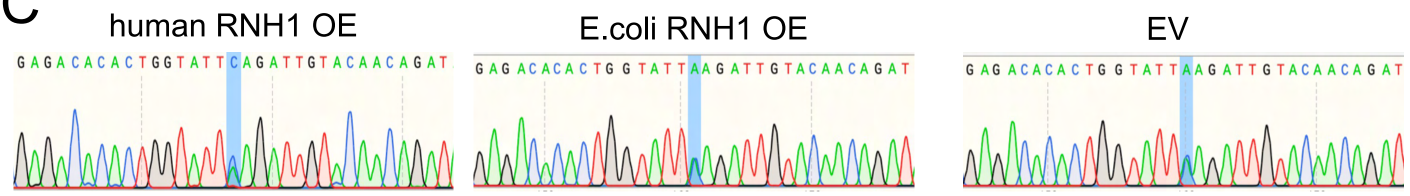
human RNH1 OE
E.coli RNH1 OE
EV

D

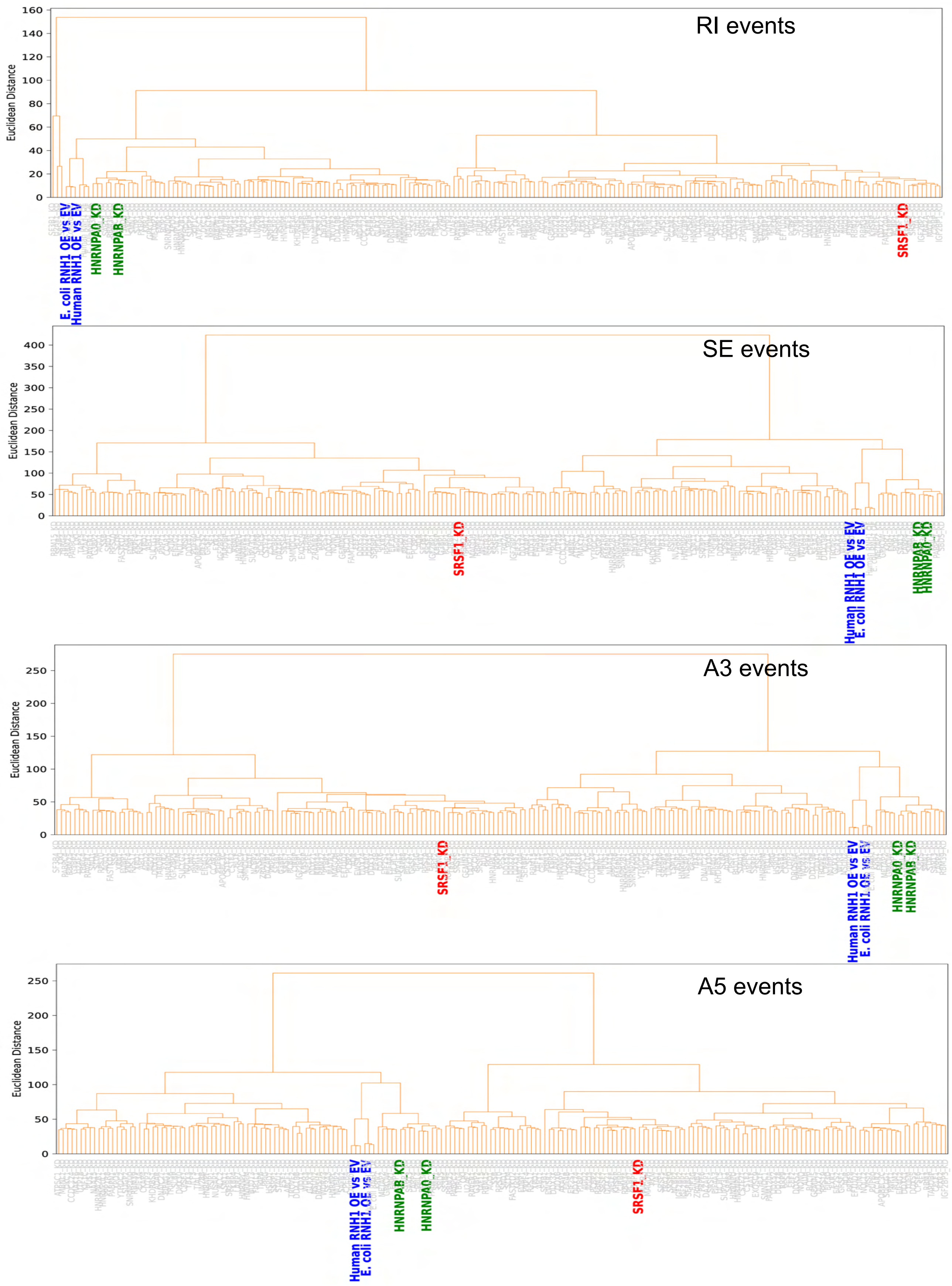
RI events
SE events
A3 events
A5 events
Euclidean Distance
E. coli RNH1 OE vs EV
Human RNH1 OE vs EV
HNRNPA0_KD
HNRNPAB_KD
SRSF1_KD
